# *SMARCA4, STK11,* and *KEAP1* co-inactivation associates with poor prognosis and upregulation of the TGF-β pathway in lung adenocarcinoma

**DOI:** 10.64898/2026.06.03.729911

**Authors:** Emily A. Costa, Barbara Pereira Mello, Corrin Wohlhieter, Subhiksha Nandakumar, Samuel Tischfield, Yingqian A. Zhan, Harsha Sridhar, Dennis Kinyua, Elisa de Stanchina, Wenfei Kang, Ning Fan, Alister Funnell, John Lazar, Justin Jee, Irina Linkov, Umeshkumar Bhanot, Esther Redin, Kristin Lee, Daniel Bates, Arielle Elkrief, Walid Chatila, Andrea Arfé, Álvaro Quintanal-Villalonga, Charles M. Rudin

## Abstract

**Background:** Lung adenocarcinoma (LUAD) is clinically and molecularly defined by oncogenic driver mutations, identification of which has led to the development of driver-targeted therapies and substantial improvements in prognosis for subsets of LUAD patients. Recent studies assessing clinical outcomes in the context of multigenic alterations have identified secondary mutations that might explain differential responses to targeted therapies, chemotherapies and immunotherapies. Genetic inactivation or loss of *SMARCA4*, which frequently co-occurs with loss-of-function mutations in *STK11* and *KEAP1*, is especially predictive of poor prognosis and shorter overall survival in LUAD patients, regardless of driver status. We sought to examine the clinical and functional associations of *SMARCA4* deficiency in LUAD, with or without co-associated *STK11/KEAP1* loss-of-function.

**Methods:** We examined correlation between *SMARCA4* loss, gene expression and prognosis through genomic and transcriptomic profiling of clinically annotated LUAD samples. We generated isogenic cell line models with genetic knockouts of *SMARCA4* with or without concomitant *STK11* and *KEAP1* to profile mutationally-defined genotypes of interest *in vitro* and *in vivo*. Lastly, we interrogated the functional dependency of *SMARCA4/STK11/KEAP1* triple mutant models on TGF-β signaling to assess its potential as a therapeutic target.

**Results:** *SMARCA4/STK11/KEAP1* triple mutant LUAD is associated with poor survival and high frequency of multisite metastasis. *SMARCA4/STK11/KEAP1* triple knockout models showed enhanced migration and invasion *in vitro*, and diversified organotropism in an *in vivo* intracardiac xenograft metastasis assay. RNA-Seq and DNaseI-Seq of these *in vitro* models and clinical samples identified upregulation of TGF-β signaling and EMT gene expression signatures, and corresponding changes in chromatin accessibility, in *SMARCA4/STK11/KEAP1* triple mutant LUAD.

**Conclusions:** We identify *SMARCA4/STK11/KEAP1* triple mutant LUAD as a prognostically significant disease subset and nominate TGF-β signaling as a potential therapeutic target.

## BACKGROUND

Lung cancer is the leading cause of cancer-related mortality globally^1^. Targeted therapies directed against oncogenic driver mutations in lung adenocarcinoma (LUAD), the most frequent lung cancer subtype, have improved patient outcomes. However, many LUAD patients lack actionable mutations and have not benefited from this progress^2^. In addition, patients who receive targeted therapy demonstrate a wide range of objective responses^3^ and almost all eventually develop resistance to these treatments^1^. Such realities fuel ongoing research to identify strategies to further improve outcomes for patients with LUAD.

LUAD is a heterogeneous disease and even canonically driver-defined subsets demonstrate diverse histological features, patterns of disease progression, and responses to targeted and other therapies. Comprehensive genomic profiling of LUAD has identified co-occurring mutations that contribute to intradriver heterogeneity^4^. Mutations in *STK11*, *KEAP1*, and *SMARCA4,* frequently observed in the context of mutant *KRAS*-driven LUAD, have been correlated with shorter survival and reduced efficacy of chemotherapy, immunotherapy, and *KRAS*^G12C^ covalent inhibitor treatment^4–10^. Recurrent inactivation of these genes has also been reported in “driverless” LUAD^2^, though their potential contributions to clinical outcomes in this setting are less understood.

Recent clinical and experimental investigation of *STK11* and *KEAP1* mutations in LUAD have yielded mechanistic insights and solidified their prognostic significance. STK11/LKB1 (serine/threonine kinase 11; liver kinase B1) is a tumor suppressor and functions as a cellular energy sensor to maintain ATP homeostasis and mitigate energetic stress^11,12^. Inactivation of *STK11* in LUAD tumor cells results in excessive proliferation and increased energetic stress, triggering metabolic dysfunction that has been explored for therapeutic targeting^13–16^. In the *Kras*^LSL(lox–stop–lox)-G12D/+^ *Trp53*^fl/fl^ (KP) genetically-engineered mouse model (GEMM) of LUAD, perturbation of *Stk11* drives tumorigenesis and acquisition of a metastatic transcriptional program through dysregulation of salt-inducible (SIK) kinases^17,18^. KEAP1 (Kelch-like ECH-associated protein 1) is a tumor suppressor that inhibits the activity of NRF2, a master regulator of antioxidant, cytoprotective, and metabolic signaling^15,19,20^. Functional loss of KEAP1 promotes tumorigenesis^15^, heightened dependence on glutaminolysis^20^, and altered glucose metabolism^19^. Dual *STK11* and *KEAP1* knockout (KO) models demonstrate heightened sensitivity to glutaminase inhibitors^21^ and pharmacological inducers of ferroptosis^22^. In the clinic, functional loss of *STK11* and *KEAP1* has been associated with increased therapeutic resistance^3,5,23–27^, higher incidence of metastasis^28^, and poor prognosis in LUAD. These associations were observed in both *KRAS*-mutant and driver-agnostic studies, suggesting universal effects of these mutations in driving aggressive, therapeutically recalcitrant LUAD^4,29^.

*SMARCA4*/BRG1 (SWI/SNF related, matrix associated, actin dependent regulator of chromatin, subfamily a, member 4; Brahma-related gene 1) is the catalytic subunit of the evolutionarily conserved SWI/SNF chromatin remodeling complex, which augments genome-wide chromatin accessibility and contributes to multiple chromatin-related processes, including transcription, repair, recombination, and replication, at tens of thousands of loci^30–32^. BRG1 (and its mutually exclusive paralog, *SMARCA2*/BRM) power both nucleosome mobilization and the release of SWI/SNF complexes from chromatin using energy derived from ATP hydrolysis^33^, and *SMARCA4* mutations that alter or disrupt its ATPase or binding abilities have been linked to oncogenic SWI/SNF activity and transcriptional dysregulation in multiple types of cancer^34–36^. In LUAD, intact *SMARCA4* acts as a tumor suppressor and inactivating mutations (which occur in 8% of LUAD) are prognostically unfavorable^8^. Molecular and functional profiling of *in vitro* and *in vivo* models of LUAD have associated *SMARCA4* loss with tumor dedifferentiation, upregulation of epithelial-mesenchymal transition (EMT) markers, and increased metastatic potential^37–39^. *SMARCA4* loss has been associated with shorter overall survival and metastatic progression in patients with LUAD^4,8,28^. However, attempts to correlate *SMARCA4* loss with specific chemotherapy and immunotherapy responses have produced conflicting findings and are likely complicated by the effects of co-mutations^6,8,40–43^.

Clinical evidence suggests that the poor outcomes associated with inactivation of *SMARCA4*, *STK11*, and *KEAP1* in LUAD may be context-dependent. *SMARCA4*, *STK11*, and *KEAP1* can be jointly affected by large deletions in chromosome 19p^4,6,41,44^. The frequency of co-loss of heterozygosity increases the likelihood that co-incident loss of these three genes occurs more often than currently recognized. Most clinical sequencing studies that have considered the mutational status of all three genes have been statistically limited by low availability of triple mutant samples. The biology underlying *SMARCA4/STK11/KEAP1* triple mutant LUAD is therefore largely uncharacterized, leaving no opportunity to leverage existing or novel therapies to improve outcomes in these patients. Here, we analyze and model *SMARCA4* loss, both with and without concurrent inactivation of both *STK11* and *KEAP1*, to deconvolute co-mutationally-defined and context-dependent impacts of *SMARCA4* deficiency in LUAD.

## MATERIALS AND METHODS

### Clinicogenomic Cohort Selection and Group Comparisons

Clinical data analyzed in this study were derived from patients at Memorial Sloan Kettering Cancer Center who underwent tumor mutational analysis using the Memorial Sloan Kettering-Integrated Molecular Profiling of Actionable Cancer Targets (MSK-IMPACT) assay^45^. Genomic alterations to *SMARCA4*, *STK11*, and *KEAP1* were filtered for oncogenic mutations using OncoKB^46^, and samples with germline alterations, variants of unknown significance, or amplifications in these genes were excluded from the analysis. All groups contain one sample per patient, prioritized by sufficient quality for FACETS analysis. If multiple samples passed quality control (QC) assessment, they were then prioritized by purity and then by the most recent sample sequenced by MSK-IMPACT. Samples in the SMARCA4 Only group have no alterations in *STK11* and *KEAP1*, and samples in the Double Mutant group have no alterations in *SMARCA4*. OncoPrints were generated in cBioPortal^47^. Demographic information for all patients whose genetic data were analyzed is in **Additional File 1, Table S1**.

### Gene and Pathway Alteration Analysis

Pathways for co-occurring and mutually exclusive gene and pathway alterations analysis were sourced from The Cancer Genome Atlas (TCGA)^48^. To enable statistical comparisons, analysis was performed only on genes with oncogenic mutations in at least 3% of the cohort. Enrichment of a co-altered gene within a group was determined by one-vs-rest, two-sided Fisher’s exact test with global p-value adjustment by FDR.

### FACETS Analysis

The FACETS algorithm and the FACETS-suite package (https://github.com/mskcc/facets-suite) were used to derive ploidy- and purity-corrected integer DNA copy-number calls, arm level events and cancer cell fraction (CCF) estimates. To ensure precise integer copy-number calls, predefined quality control metrics were applied (https://github.com/taylor-lab/facets-preview). Samples that didn’t pass QC metrics were excluded. Biallelic inactivation was defined as either homozygous deletion, two or more deleterious/oncogenic mutations in the same tumor sample, one mutation with concurrent heterozygous loss of the wild-type (WT) allele, or one mutation with concurrent copy-neutral loss-of-heterozygosity (CN-LOH) as computed using the FACETS algorithm^49^. Sample numbers for this analysis are listed in **Additional File 1, Table S2**.

### Overall Survival and Hazard Ratio Analysis

Kaplan–Meier curves for whole cohort survival were constructed for overall survival (OS) and compared via the Mantel–Cox log-rank test. To assess associations between clinical and genomic features and OS, we estimated each hazard ratio (HR) using Cox proportional hazards regression models. Multiple testing correction was performed using the FDR method (adjusted p-value cutoff of 0.05). All analyses were performed using R v4.3.1.

### Metastasis Incidence and Site Frequency Analysis

Metastatic site data extraction was performed by the MSKCC Clinical Data Mining Team. For patients with available metastatic site data, sites were extracted via Natural Language Processing from radiology reports or sourced metastasis site annotations in cBioPortal. Metastatic sites were categorized as: adrenal glands, intra-abdominal, bone, central nervous system (CNS)/brain, liver, pleura, reproductive organs, and other. Lymph nodes and secondary lung metastases were omitted. For radiology report data extraction, a ClinicalBERT transformer model^50^ was fine-tuned in a multi-labeling setting, so that multiple positive sites of disease may be predicted for a single report. The network was trained to jointly predict disease across all sites. During training, the network was trained to minimize the sum of 10 per-site binary cross-entropy losses from individual per-site binary-prediction logits. Pairwise comparisons were done by two-sided Fisher exact test. Sample numbers for this analysis are listed in **Additional File 1, Table S3**.

### Chemotherapy and Immunotherapy Survival Analyses

Patients in the chemotherapy cohort received any cytotoxic chemotherapy (carboplatin, pemetrexed, cisplatin, paclitaxel, gemcitabine, docetaxel, vinorelbine, etoposide) without concurrent immunotherapy. Patients in the immunotherapy cohort received any checkpoint inhibitor (durvalumab, ipilimumab, nivolumab, atezolizumab, pembrolizumab) with or without concurrent chemotherapy. Medication data were obtained from structured institutional medication and pharmacy records, and real-world patient data were derived as described^51^. Kaplan-Meier curves were generated from the time of first initiation of therapy class (chemotherapy or immunotherapy) to the beginning of the next treatment regimen or death, right censored at the date of last follow-up. P-values were calculated using Cox proportional hazards models using Python 3.10 and the lifelines 0.27.7 package. Sample numbers for this analysis are listed in **Additional File 1, Table S4**.

### Clinical Sample Selection

Surgically resected LUAD tumor tissue samples were sourced from the MSKCC Precision Pathology Biobanking core facility. All samples were derived from the primary tumor site, had annotated oncogenic mutations in *SMARCA4*, *STK11*, and *KEAP1*, lacked mutations to other SWI/SNF complex members screened in MSK-IMPACT, and were flash frozen within 4 hours of resection to enable high-quality RNA-Seq. Clinical demographics and characteristics of the patients these samples came from are in **Additional File 1, Tables S5 and S6**. Methods for sample preparation and RNA sequencing are in **Supplementary Methods.**

### Cell Lines and Culturing Conditions

NCI-H358 (ATCC, catalog no. CRL-1848) cells were cultured in RPMI media supplemented with 10% FBS and 1X Penicillin-Streptomycin (PS) and incubated in 5% CO2 at 37° C. Their identity was confirmed by STR profiling before and after single cell cloning and cultures were periodically verified as mycoplasma-negative using a MycoAlert® Mycoplasma Detection Kit (Lonza, catalog no. LT07-318). Methods for single cell cloning are in **Supplementary Methods.**

### CRISPR/Cas9 Genetic Engineering

The cell lines were transduced with lentivirus containing lentiCas9-Blast (Addgene, catalog no. 52962). *STK11/KEAP* double knockout (DKO) cell line engineering is described^22^. Candidate *SMARCA4* knockout guide RNA (sgRNA) sequences were generated by GuideScan^52^ and the 5 guides with the highest cutting efficiency and specificity scores and fewest off targets were selected for testing. Candidate guide oligos and a safe-targeting sgRNA (sgSAFE) were annealed and cloned into lentiGuide-Neo (Addgene, catalog no. 139449) using single-step digestion-ligation with BsmBI. Lentiviruses for each *SMARCA4* sgRNA plasmid were transduced at a high MOI to 293T cells and knockout was assessed by Western blot and Tracking of Indels by Decomposition (TIDE)^53^. The *SMARCA4* sgRNA that resulted in the greatest decrease in BRG1 expression was selected for SMARCA4^KO^ cell line engineering. sgSMARCA4 lentiGuide-Neo was transduced to H358 cells, and KO cells were selected by neomycin treatment (200ug/mL). All sgRNA sequences are listed in **Additional File 1, Table S8** and methods for lentiviral production and transduction are in **Supplementary Methods.**

### RNA-Seq Gene Differential Expression Analysis

Raw FASTQ files from RNA-seq paired-end sequencing were aligned to the ENSEMBL GRCh37 Homo sapiens release 99 transcriptomes using Kallisto^54^. Gene expression levels (calculated as transcripts per million) and differential expression was performed using Sleuth (v0.30.0)^55^. Sleuth was used to plot PCA and the percentage of variance explained based on Transcripts Per Million (TPM) using the functions “plot_pca” and “plot_pc_variance”. Heatmaps of gene expression show the Z-score (scaled by rows to normalize across cell lines) of TPM. The heatmap in **Figure 4D** was made in SRplot. Differentially expressed genes across the genotypes were identified using the following model: Expression ∼ MutationStatus. Pathway analysis was performed in R using the package clusterProfiler (https://bioconductor.org/packages/release/bioc/html/clusterProfiler.html) with three gene set libraries, mSigDB, KEGG and GO. Gene ranking was calculated by taking −log10(pval)*sign(b)) and pathways with p-value > 0.05 were culled. Methods for sample preparation and RNA sequencing are in **Supplementary Methods.**

### Western Blots

Whole cell lysates were extracted from frozen cell pellets, which were resuspended in ice cold RIPA buffer (Pierce, catalog no. 89901) and 1X Halt^TM^ protease and phosphatase inhibitor (Thermo Scientific, catalog no. 78446), incubated on ice for 20 minutes, sonicated for 10 seconds at 40% amplitude using a Model 120 Sonic Dismembrator (Fisher Scientific), and centrifuged at 13.6 x g for 30 minutes at 4° C. Supernatant was collected and total protein was quantified by Pierce^TM^ BCA protein assay (Thermo Scientific, catalog no. 23227). Lysates were diluted to the same concentration in NuPAGE^TM^ LDS sample buffer (Invitrogen, catalog no. NP0007) and NuPAGE^TM^ sample reducing agent (Invitrogen, catalog no. NP0009) and heated at 75° C for 10 minutes. 30-60μg of total protein was loaded to each well of a precast NuPAGE^TM^ 4-12% Bis-Tris Gel (Invitrogen) and run at 50V through the resolving phase and 120V until completed. Proteins were wet transferred to a 0.45µM Immobilon-P PVDF membrane (EMD Millipore, catalog no. IPVH00010) overnight in a solution of 1X Tris-Glycine with 20% methanol. Blots were blocked in a 5% non-fat dry milk in TBS buffer and 0.1% Tween-20 (TBS-T) solution with gentle shaking for one hour at RT or overnight at 4° C. Following TBS-T rinsing, blots were probed with primary antibodies prepared at a dilution of 1:1000 in 5% bovine serum albumin (BSA) or 5% milk in TBS-T overnight at 4° C with gentle shaking. Blots were then washed with vigorous shaking 4 times (10 minutes each) with TBS-T, probed with secondary antibody prepared at a dilution of 1:3000 in TBS-T or 5% milk in TBS-T for one hour, and washed 4 times in TBS-T as previous. Protein expression was detected via ECL with Pierce^TM^ ECL Western Blotting Substrate (Thermo Scientific, catalog no. 32209) or SuperSignal^TM^ West Pico PLUS Chemiluminescent Substrate (Thermo Scientific, catalog no. 34577) and imaged on an iBright^TM^ CL1500 Imaging System (Invitrogen). All primary and secondary antibodies are listed in **Additional File 1, Table S7**. All Westerns were performed in triplicate from 3 separately collected and prepared sets of lysates to verify reproducibility. Data shown in figures are representative results.

### BRG1 Western Blot Expression Quantification

BRG1 expression was quantified via pixel density measurement in Fiji^56^. ROIs of equal dimensions were used to measure each band as well as capture background measurements from space just above or below each band. Band measurements were inverted (255-X) and background subtracted, and then BRG1 expression for each lane was calculated as a ratio of BRG1 band pixel density over loading control (cyclophilin B) band pixel density. BRG1 expression in *SMARCA4* knockout cell line clones is shown as percentage of mean BRG1 expression in sgSAFE cell line clones, with mean BRG1 expression in the sgSAFE clones set equal to 100%.

### TGF-β1 ELISA

Two days before seeding the assay, equal numbers of clones of the same KO genotype were pooled and seeded. 25,000 cells were seeded in quadruplicate to a 96-well white microplate in 150μL serum-free media and cultured for 96 hours. TGF-β1 standard curves and immunoassay control groups (Quantikine Immunoassay Control Group 1, biotechne, catalog no. QC01-1) were prepared in triplicate and according to the manufacturer’s protocol (Human/Mouse/Rat/Porcine/Canine TGF-beta 1 Quantikine ELISA, biotechne, catalog no. DB100C). TGF-β1 was activated and assayed as described in the protocol, and cell culture supernate samples were tested neat so the dilution factor was 1.4. Following addition of the detection reagents and required incubation periods, the optical density of all wells (assay wells, standard curve wells, and control group wells) were read on a BioTek Synergy Neo microplate reader (Agilent) at 450nm and 570nm. Reads at 570nm were subtracted from reads at 450nm to correct for plate imperfections. Read triplicates and quadruplicates were averaged and multiplied by 1.4, and the standard curve was generated using a four-parameter logistic fit. Concentrations of TGF-β1 in assay wells were ascertained by interpolation from the standard curve.

### Cell Morphology Microscope Imaging

Cells were seeded to 6-well plates at a low seeding density and allowed to adhere and grow for 3 to 4 days prior to imaging (bright field, 20X magnification 20x/0.8NA objective) on a Zeiss Axio Observer.Z1 microscope.

### Migration Assay

Two days before serum starving, equal numbers of clones of the same KO genotype were pooled and seeded (e.g. SAFE clone were combined, DKO clones were combined, etc.). Cells were placed in FBS-free RPMI media and serum starved 24 hours before initiation of the assay. To start the assay, serum-starved cells were trypsinized, counted, and reconstituted in no FBS media. 150,000 cells per insert were seeded to transwell inserts (Falcon Permeable Support, Corning, catalog no.353182). Experimental inserts were seeded in triplicate (with FBS-containing media in the receiver well) with one control insert for each cell line (with no FBS media in the receiver well). Simultaneously, 150,000 cells from the same cell mixture that was seeded to the inserts was seeded in triplicate to a 24-well plate, which was fixed with 4% paraformaldehyde and Crystal Violet with 20% methanol the following morning to confirm equal seeding. Cells in the migration assay were incubated for 48 hours after seeding. To complete the assay, the transwell inserts were carefully lifted from the receiver plates with forceps, the inside of the insert was swabbed with a media dampened cotton swab to remove non-migrated cells, and the bottom surface of the insert was fixed first in 4% paraformaldehyde for 15 minutes and then in a solution of Crystal Violet and 20% methanol for 15 minutes before being rinsed with water and allowed to dry. Crystal Violet stained (migrated) cells on each insert were imaged on an EVOS FL microscope (bright field, 20X magnification) for quantification. 5 fields were captured per insert. Cells were counted using the Cell Counter tool in Fiji. The reported number of migrated cells is the sum of experimental insert cells that migrated in 5 fields subtracted by the number of cells that migrated through the control insert. This assay was performed three times. Data shown in figures are representative results from one assay.

### Invasion Assay

Two days before serum starving, like clones were pooled in equal numbers and seeded. Cells were placed in RPMI media without FBS (no FBS media) and serum starved 24 hours before initiation of the assay. Prior to seeding the assay, BioCoat^TM^ Matrigel-coated transwell inserts (Corning, catalog no. 354480) were thawed at room temperature and rehydrated in warm no FBS RPMI media for 2 hours at 37° C. Cells were seeded at 200,000 cells per insert and incubated for 72 hours. Inserts were swabbed, fixed, stained, and imaged as described for the migration assays. A control plate with cells seeded from the assay mixtures and fixed the morning after the assay was seeded was also prepared as previously described and assessed for equal seeding. Cells were quantified as previously described. This assay was performed three times. Data shown in figures are representative results from one assay.

### Intracardiac Xenograft Injections and Bioluminescence Imaging (BLI)

The xenograft assays took place in mice (NOD.Cg-Prkdcscid Il2rgtm1Wjl/SzJ (NSG) females, 6-8 weeks old, Charles River Laboratories) in accredited facilities and under pathogen-free conditions. Luciferase-expressing single clone cell lines of the same genotype were mixed in equal numbers at most two days before injection. Cells were administered via intracardiac injection (10,000 cells per injection) to 10 mice per group at T0. One mouse in the SAFE group passed shortly after intracardiac injection, so n=9 for this group. Mice were injected with D-Luciferin and imaged weekly by bioluminescence imaging (BLI) by IVIS (Perkin Elmer) to monitor tumor growth. This animal experiment was approved by the Memorial Sloan Kettering Cancer Center (MSKCC) Animal Care and Use Committee.

### IVIS Signal Quantification and Tumor Growth Analysis

IVIS BLI signal were measured in Living Image software (PerkinElmer). Rectangular Measurement ROIs were drawn over each mouse (from nose to the base of the tail) from a dorsal view to capture Average Radiance and standard deviation. Signal over time is plotted starting at the first imaged time point after injection. A general least squares (GLS) approach was used to test for differences in temporal trajectories in log-average radiance across time points. Accordingly, a general unspecified covariance structure was assumed to account for the potential within-mouse correlation across each time-point. Parameters were estimated by the maximum likelihood method, and tests of fixed effects implemented using likelihood ration statistics. Analyses were performed using the “nlme” (v 3.1-166) R package.

### *In vivo* Metastasis Quantification

Slide images were randomized and blinded prior to image quantification by two methods: counting tumor foci by eye and measuring tumor area by semi-supervised neural network-based area estimation in QuPath^57^. A script was written in QuPath to automate the measurement of tumor and non-tumor areas in each sample using the pixel classifier tool, and measurements were confirmed visually to ensure accurate classification. Quantification was performed by the same person for all images and the images blinded during quantification. Methods for necropsy, sample preparation, and immunohistochemistry are in **Supplementary Methods.**

## RESULTS

### *SMARCA4* only mutant and *SMARCA4/STK11/KEAP1* triple mutant tumors are characterized by distinct co-mutational patterns

To better define the clinical implications of *SMARCA4* loss, with or without co-loss of both *STK11* and *KEAP1*, we performed genomic analyses of LUAD tumors. Among all LUAD tumors sequenced by MSK-IMPACT, mutations in *SMARCA4*, *STK11*, and *KEAP1* were present at frequencies of 8%, 16%, and 14%, respectively **(Supplementary Figure S1A)**. As this study focuses on context-specific biology of *SMARCA4*, we selected tumors with loss-of-function mutations in *SMARCA4*, *STK11*, and *KEAP1* but excluded tumors with *STK11* only, *KEAP1* only, *STK11/SMARCA4* only, and *KEAP1/SMARCA4* only mutations. The resulting tumor sequencing cohort (n=5048) contained four mutationally-defined groups: tumors with mutations in *SMARCA4* (“SMARCA4 Only” group, n=129); *STK11* and *KEAP1* (“Double Mutant” group, n=180); *SMARCA4*, *STK11*, and *KEAP1* (“Triple Mutant” group, n=47); and tumors in which all three genes were wild-type (“Unaltered” group, n=4692). There were no overlapping samples or patients between the groups **(Figure 1A)**.

**Figure 1.**
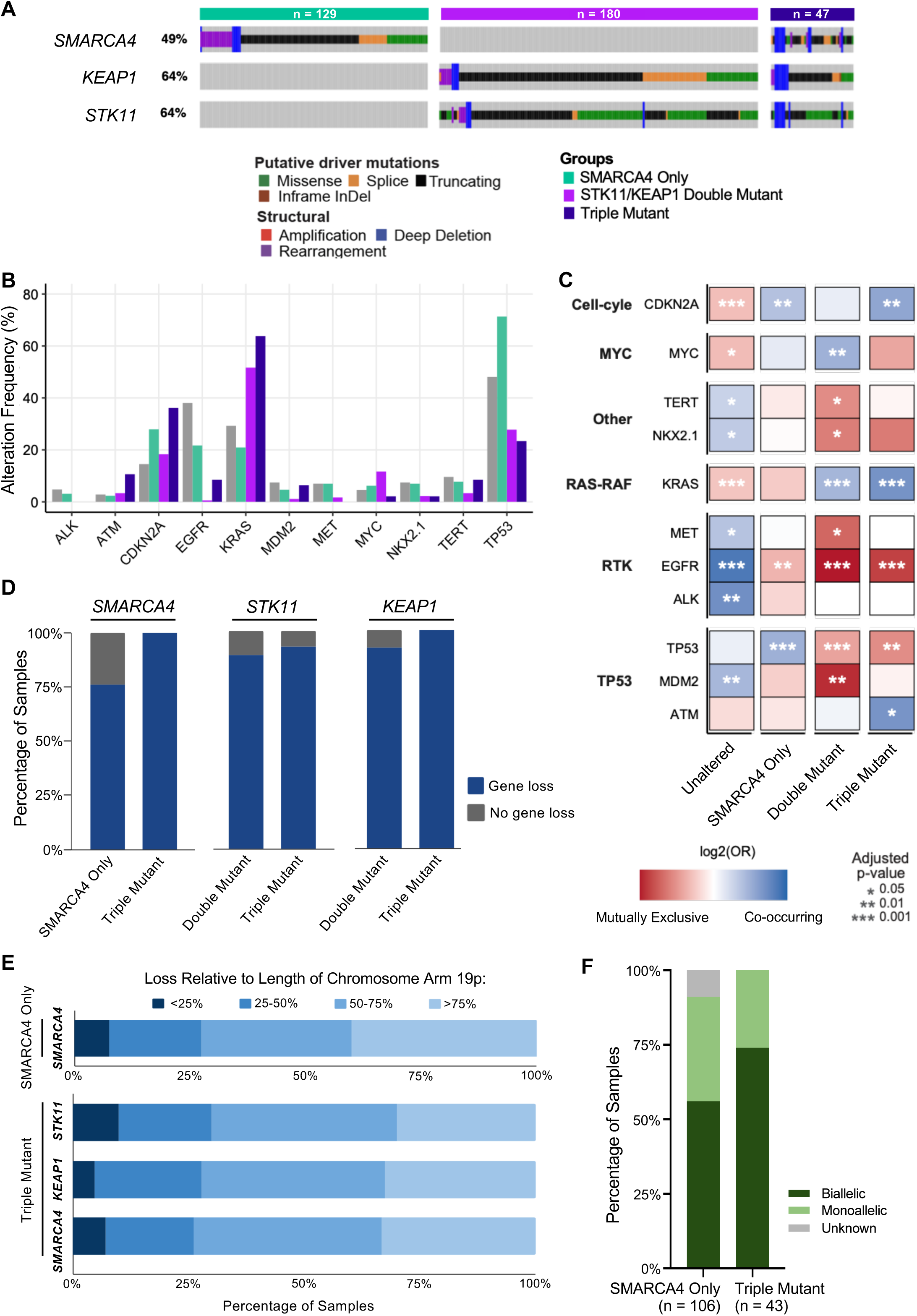
Triple Mutant tumors have distinct co-mutational patterns and universal loss of *SMARCA4* and *KEAP1* genes. **A.** MSK-IMPACT OncoPrint of *SMARCA4*, *STK11*, and *KEAP1* genetic alterations in the clinicogenomic tumor sequencing cohort. Each vertical line is a single sample. There is no sample overlap among the SMARCA4 Only, Double Mutant, and Triple Mutant groups. **B.** Bar plot showing alteration frequencies of selected genes co-altered in ≥ 3% of the cohort. **C.** Enrichment analysis of co-altered genes. Genes are grouped by shared pathway annotations. OR = Odds Ratio. Enrichment of a co-altered gene within a group was determined by one-vs-rest, two-sided Fisher’s exact test with global p-value adjustment by FDR. *p<0.05, **p<0.01, ***p<0.001. **D.** Stacked bar plots of the percentages of samples with gene loss in *SMARCA4, STK11,* and *KEAP1* as defined by the FACETS algorithm (homozygous or heterozygous chromosomal deletions). **E.** Stacked bar plots of the percentages of samples per group with the shown amounts of chromosome 19p arm loss (<25%, 25-50%, 50-75%, >75%). Focal deletions are defined as <25% arm loss. Broad, arm-level deletion events are defined as >50% arm loss. **F.** Stacked bar plots showing the percentages of samples per group with monoallelic or biallelic loss of *SMARCA4*. Samples in gray could not be assessed by FACETS.

We observed significantly more patients with a smoking history **(Supplementary Figure S1B)** and increased tumor mutational burden (TMB) **(Supplementary Figure S1C)** in the SMARCA4 Only, Double Mutant, and Triple Mutant groups compared to the Unaltered group. These results align with previous LUAD studies showing that mutations in *SMARCA4*, *STK11*, and *KEAP1* individually correlate with a history of smoking^8,29^ and that *SMARCA4* deficiency correlates with higher TMB^8,41^. Adding to these observations, our data indicate that, in the context of *STK11/KEAP1-*mutant LUAD*, SMARCA4* mutations do not substantially impact the association of *STK11* and *KEAP1* mutations with smoking history or TMB.

Next, we sought to compare co-occurrence and mutual exclusion patterns of other known cancer-related genes across our four comparator groups. To enable statistical comparisons, our co-alteration enrichment analysis only included genes with oncogenic mutations in at least 3% of the cohort. We observed a much stronger association with TP53 mutations in the SMARCA4 Only group than in the other groups (p<0.001), while, conversely, KRAS mutations are more prevalent in the Double and Triple Mutant groups (p<0.001) **(Figure 1B and C)**. Contrasting reports from other *SMARCA4*-focused sequencing studies, we did not observe enrichment for co-occurring *KRAS* mutations in our SMARCA4 Only group^6,8^. Notably, these studies did not (and lacked the necessary sample sizes to) analyze co-mutation of *KRAS* and *SMARCA4* in the absence of co-altered *STK11* and *KEAP1*. We captured previously reported gene co-alterations including mutual exclusivity with *EGFR* mutations in the SMARCA4 Only, Double Mutant, and Triple Mutant groups (SMARCA4 Only: p<0.01, Double and Triple Mutant: p<0.001)^4,8,28,58^. Enrichment for co-occurrence of *ATM* mutations was also observed in the Triple Mutant group (p<0.05).

As *SMARCA4*, *STK11*, and *KEAP1* inactivation frequently involves loss or co-loss of heterozygosity (LOH) events, we were also interested to see whether genetic inactivation of *SMARCA4* differed between the SMARCA4 Only group and the Triple Mutant group. Homozygous inactivation of *SMARCA4* has been estimated to occur in 70-90% of *SMARCA4*-mutant NSCLC^41^ and in most cases (∼80%) involves loss of heterozygosity^6,41^. Of the remaining intragenic SMARCA4 mutations, about half are missense mutations while the other half are truncating mutations^6,8^, the latter of which more often co-occur with co-loss of *STK11* and *KEAP1* and correlate with lower survival in metastatic NSCLC patients^8,41^. As observed previously^6^, *SMARCA4* mutations in our cohort were diffuse across the gene body, with no apparent selection for functional domains **(Supplementary Figure S1D)**. We also observed no major differences in their distribution between the SMARCA4 Only and Triple Mutant groups.

Gene loss, i.e. complete loss of at least one allele (inclusive of focal deletions, arm-level events, and either heterozygous or homozygous loss), was noted in *SMARCA4*, *STK11*, or *KEAP1* in 85% of the samples across the SMARCA4 Only, Double Mutant, and Triple Mutant groups **(Figure 1D)**. All samples in the Triple Mutant Group had loss in *KEAP1* and *SMARCA4*. In the SMARCA4 Only and Triple Mutant groups, >50% of losses in all three genes were broad, arm-level deletions, affecting >50% of the chromosome arm. In the Triple Mutant group, *STK11* displayed the greatest propensity for focal events while *SMARCA4* and *KEAP1* deletion patterns were nearly identical **(Figure 1E)**. These phenomena, particularly the overlapping patterns of *KEAP1* and *SMARCA4* loss in the Triple Mutant group, likely stem from the relative positioning of the three genes on chromosome 19p—*STK11* is about 9 Mb away from *SMARCA4* and *KEAP1*, which are themselves separated by about 0.5 Mb^59^. Comparing the incidence of monoallelic versus biallelic (or heterozygous versus homozygous) loss of *SMARCA4*, we observed that *SMARCA4* biallelic loss was numerically more frequent in the Triple Mutant versus SMARCA4 Only group (74% vs. 56%), though this apparent difference did not reach statistical significance **(Figure 1F)**. Irrespective of *STK11/KEAP1* co-mutations, gene deletion commonly contributes to *SMARCA4* inactivation in LUAD, with arm-level deletions on chromosome 19p driving the high frequency of co-LOH with *STK11* and *KEAP1*.

### *SMARCA4/STK11/KEAP1* triple mutant patients have the lowest survival and highest incidence of metastasis to multiple sites

We next sought to define the prognostic effects of *SMARCA4* loss with and without co-concurrent loss of *STK11* and *KEAP1*. In our cohort, the Triple Mutant group displayed the shortest overall survival (HR=4.5 vs. Unaltered group, p<0.001), with a hazard ratio about 3-fold greater than those of the SMARCA4 Only and Double Mutant groups (HR=3.03 vs. SMARCA4 Only, p<0.001; HR=2.67 vs. Double Mutant, p<0.001) **(Figure 2A; Supplementary Figure S2A)**. The Triple Mutant group also conferred a higher risk for shorter overall survival than oncogenic *KRAS* or a history of smoking and a relative risk numerically higher than Stage 4 versus Stage 1-3 diagnosis **(Figure 2B)**.

**Figure 2.**
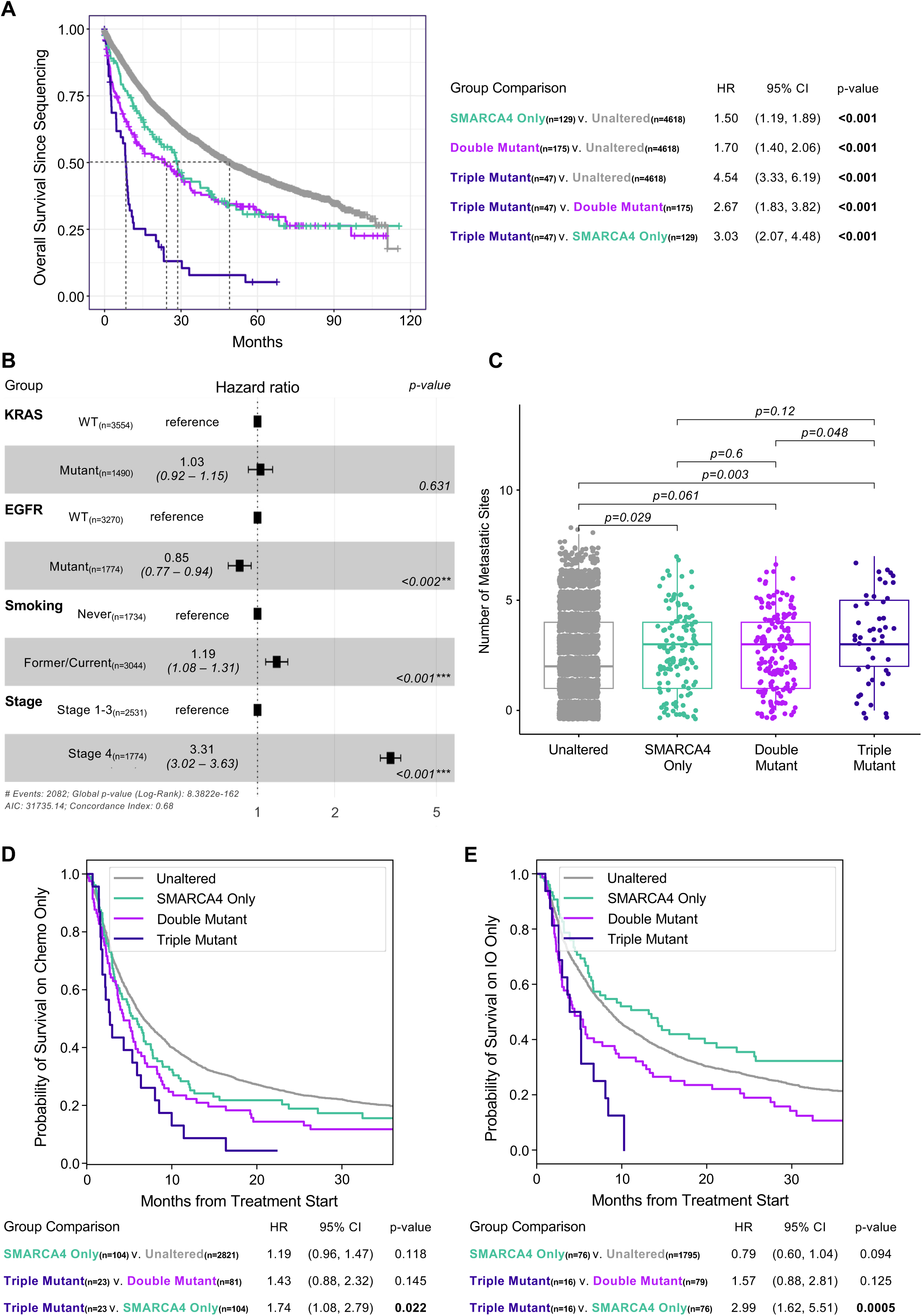
Triple Mutant patients have the lowest survival and highest incidence of metastasis to multiple sites. **A.** Kaplan-Meier curves of overall survival for each cohort group, with hazard ratios (HR) estimated using Cox proportional hazards regression models. Dotted lines show median survival. CI = Confidence Interval. See also Supplementary Figure S2A. **B.** Table of hazard ratios of overall survival per co-variate (*KRAS and EGFR* status, smoking history, and disease stage) over reference, estimated as in Figure 2A. WT = wild-type. **C.** Plotted metastasis sites per patient (0 to 7 sites) across groups. The center line is the median score. Pairwise comparisons were done by two-sided Fisher exact test. P-values are shown above brackets. **D.** Chemotherapy (chemo) cohort and **E.** immunotherapy (IO) cohort Kaplan-Meier curves of overall survival, with hazard ratios (HR) estimated as in Figure 2A. CI = Confidence Interval. See also Supplementary Figure S2C.

Individual genetic inactivation of *Smarca4*, *Stk11*, and *Keap1* has been shown to promote tumor progression and metastasis in the KP LUAD GEMM^20,37,60,61^. Recent clinical profiling of LUAD primary and metastatic tumor samples noted correlation of *SMARCA4* and *KEAP1* alterations and high frequencies of *SMARCA4/STK11* and *SMARCA4/KEAP1* co-alteration with ever-metastatic primary tumors, and of *SMARCA4* alterations with a shorter time to metastasis^28^, with another study extending these observations to KRAS-mutant metastatic LUAD^62^. We therefore speculated that differential incidence of metastasis among our cohort groups might be a major contributor to their range in overall survival.

To explore this, we assessed sites of extrapulmonary metastasis from the patients in our cohort from radiology reports. Despite substantial intergroup heterogeneity in the extent of metastasis, a statistically significant and greater number of metastatic sites were observed in patients in the SMARCA4 Only versus Unaltered group (p=0.029) and in the Triple Mutant versus Double Mutant group (p=0.048), consistent with *SMARCA4* loss promoting metastatic spread **(Figure 2C)**. While not reaching statistical significance, the Triple Mutant group had the highest percentage of patients with over five sites of metastasis and the highest frequencies of metastasis in both canonical LUAD metastatic sites (i.e. adrenal glands, brain, and bone^28^) and in more unusual sites (intra-abdominal and “other”) **(Supplementary Figure S2B and C)**.

We were interested to compare survival on standard courses of therapy across our cohort groups. *STK11* and *KEAP1* inactivating mutations are strongly associated with resistance to chemotherapy and immune checkpoint inhibitor (ICI) therapy^10,24,27,29,63–65^, particularly when they co-occur with oncogenic *KRAS*^25^. However, as with studies reporting overall survival, many retrospective studies correlating *SMARCA4* status with therapy response did not account for the potential prognostic impact of *STK11* and *KEAP1* status. We identified patients from our cohort groups who had received only cytotoxic chemotherapy or checkpoint inhibitor therapy and performed therapy-focused overall survival analyses **(Figure 2D and E; Supplementary Figure S2D and E)**. While these subset analyses lose statistical power for some comparisons, in both the chemotherapy and immunotherapy analyses, survival in the Triple Mutant group was significantly lower than in the SMARCA4 Only group (p=0.022 and p=0.0005, respectively), indicating that the addition of *STK11/KEAP1* inactivation is prognostically impactful in the context of in *SMARCA4-*mutant LUAD. The Triple Mutant group had the worst outcome on both chemotherapy and immunotherapy. Surprisingly, the SMARCA4 Only group showed (statistically insignificant) improved survival over all others in the immunotherapy analysis (HR=0.79 vs. Unaltered group), warranting further investigation of immunotherapy efficacy in a larger population of these patients.

These findings indicate that patients with *SMARCA4*/*STK11/KEAP1* triple mutant LUAD have the worst prognosis and tend to have greater metastatic burden than others. These data further suggest that *STK11/KEAP1* context might underlie previously conflicting reports of therapy responses in *SMARCA4*-deficient LUAD patients, underscoring the imperative to consider co-incident alterations in *SMARCA4, STK11,* and *KEAP1* as factors affecting LUAD patient outcomes.

### *SMARCA4* loss gene signatures vary with *STK11/KEAP1* co-mutational status

To characterize the transcriptional profiles associated with *SMARCA4* loss with or without *STK11/KEAP1* loss, we performed RNA-Seq on a curated set of LUAD clinical samples. These samples were exclusively derived primary lung tumors, eliminating potential variability imposed by distant organ microenvironments. This RNA-Seq cohort included 11 SMARCA4 Only, 11 Double Mutant, 9 Triple Mutant, and 18 Unaltered cases **(Supplementary Figure S3A)**. In addition to screening for inactivating mutations in *SMARCA4*, all tumors were stained for BRG1 protein expression. One *STK11/KEAP1* mutant tumor had intact *SMARCA4* by sequencing but entirely lacked detectable BRG1 protein and was therefore included in the Triple Mutant group.

The RNA-Seq cohort generally reflected clinicogenomic observations made in the larger MSK-IMPACT cohort. The most frequently mutated driver oncogene among the samples was *KRAS* (51%) **(Supplementary Figure S3A)**. Frequencies of *KRAS* alteration in the Double and Triple Mutant groups were similar to the MSK-IMPACT cohort **(Figure 1B)**, while the SMARCA4 Only group had more frequent *KRAS* alterations. Additionally, the Triple Mutant group of the RNA-Seq cohort had the highest proportion of patients with metastasis, the highest proportion of patients with multiple metastatic sites, and the broadest variety of metastatic sites **(Supplementary Figure S3B-D)**.

To assess gene expression changes specifically associated with *SMARCA4* loss, we compared the SMARCA4 Only versus Unaltered and Triple Mutant versus Double Mutant groups. Differential gene expression (DGE) and pathway enrichment analyses of both comparisons revealed dysregulation of genes in pathways involved in malignant transformation, including cell cycling, metabolic stress, developmental signaling, and EMT. Previously reported transcriptional effects of *SMARCA4* loss were confirmed in our data. Upregulation of pathways linked to mitochondrial physiology and oxidative phosphorylation (OXPHOS) aligns with elevated OXPHOS-associated gene expression and inhibitor sensitivity observed in *in vivo* models of *SMARCA4*-deficient LUAD^37,66^ **(Figure 3A; Additional File 2, Tables S1-4)**. Pathways related to cell cycle progression were also upregulated, consistent with the known interaction of BRG1 with pRB, inhibiting transcription of E2F genes and suppressing G1→S checkpoint transition^67^ **(Figure 3B; Additional File 3, Tables S1-4)**.

**Figure 3.**
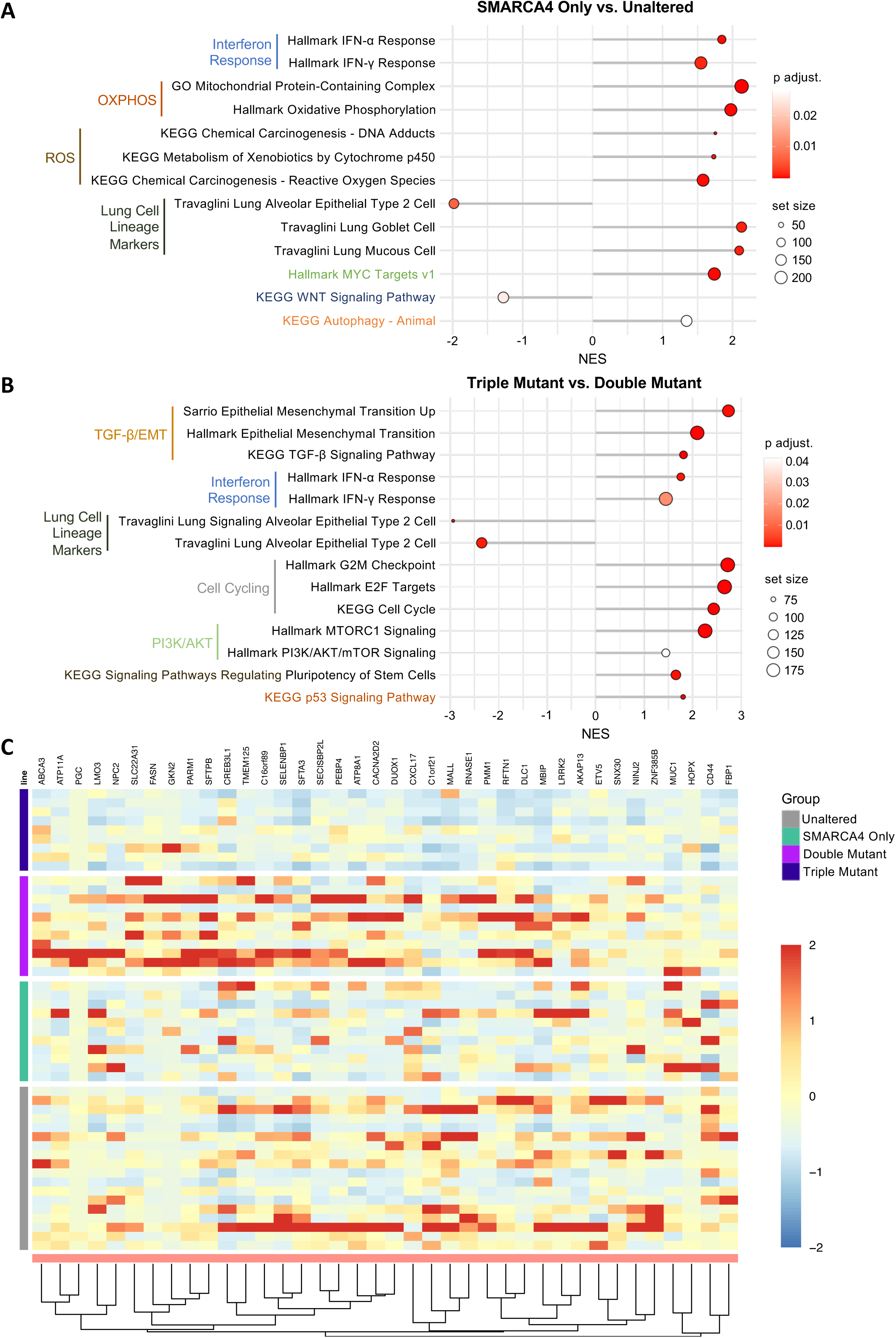
Differential *SMARCA4* loss-associated gene signatures appear with and without *STK11/KEAP1* co-mutation in clinical samples. Gene Set Enrichment Analysis (GSEA) of differentially expressed genes (DEGs) between the **A.** SMARCA4 Only vs. Unaltered clinical sample groups and **B.** Triple Mutant vs. Double Mutant clinical sample groups. All pathways shown are significantly enriched (adjusted p<0.05) and from the KEGG, GO, and mSigDB gene set databases. NES = normalized enrichment score. See also Additional Files 2 and 3. **C.** Heat map showing relative expression of leading edge genes from the “Travaglini Lung Signaling Alveolar Epithelial Type 2 Cell” gene set in Triple Mutant vs. Double Mutant clinical sample GSEA. Each column in the heatmap is a single clinical sample. Hierarchical clustering was performed using Euclidean distance and the “complete” clustering method. Shading corresponds to Z-score of gene transcripts per million (TPM) normalized by row.

Our analysis additionally revealed unique transcriptional associations with *SMARCA4* deficiency regardless of *STK11/KEAP1* status, such as upregulation of interferon (IFN)-α and -γ signaling genes, which have been previously associated with acquired immunotherapy resistance in LUAD^68^. We also observed downregulation of alveolar type II (AT2) cell lineage markers **(Figure 3A and B**; Additional File 2, Tables S1-4; Additional File 3, Tables S1-4). Histologically, *SMARCA4*-deficient lung cancers are characterized by undifferentiated morphology and frequent loss of epithelial differentiation markers^69–71^, and loss of AT2 and club cell lineage markers has been specifically observed in *SMARCA4* deficient LUAD^37^. In our RNA-Seq data, loss of AT2 lineage expression in the SMARCA4 Only versus Unaltered group comparison was accompanied by upregulation of pathways associated with mucin-producing (mucinous) cells **(Figure 3A; Additional File 2, Tables S1-4)**. Across all four groups, AT2 markers were most downregulated in the Triple Mutant samples, which uniformly demonstrated low expression of these genes **(Figure 3C)**. These findings align with previous histological assessment of *SMARCA4*-mutant LUAD and further suggest that triple mutant LUAD is less differentiated than *SMARCA4*-only mutant LUAD.

A strong transcriptional signature that emerged from the Triple Mutant versus Double Mutant group comparison was the upregulation of TGF-β signaling and EMT-associated genes **(Figure 3B; Additional File 3, Tables S1-4)**. This finding, not seen in SMARCA4 Only versus Unaltered group comparison, is of particular interest considering the exceptionally poor outcomes and high metastatic burden we observed in patients with triple mutant LUAD, as TGF-β signaling is well-known to support lung cancer metastasis through induction of EMT^72^.

These results identify transcriptional signatures associated with *SMARCA4* mutation in LUAD primary tumors, capture effects of differential *STK11/KEAP1* status, and nominate upregulation of TGF-β signaling as a link between the biology of *SMARCA4/STK11/KEAP1* triple mutant tumors and their observed propensity for metastasis and poor clinical outcomes.

### *SMARCA4/STK11/KEAP1* co-inactivation promotes TGF-β signaling and an EMT phenotype

To investigate the molecular basis of our clinicogenomic and transcriptomic observations, we used CRISPR/Cas9 to genetically engineer isogenic single cell clone models with knockout (KO) of *SMARCA4*, *STK11*, and *KEAP1* in the NCI-H358 LUAD cell line, which has an oncogenic *KRAS^G12D^* mutation. Mimicking our clinical cohorts, we generated isogenic lines with the following combinations of stable gene knockouts: *SMARCA4* only knockout (SMARCA4^KO^), *STK11* and *KEAP1* double knockout (DKO, as previously reported^22^), triple knockout of all three genes (TKO), and a control knockout with a safe-targeting guide RNA (SAFE). Knockouts were confirmed by Western blotting for BRG1, LKB1, and KEAP1 **(Figure 4A)**. BRM expression slightly increased in most SMARCA4^KO^ clones and, as expected, NRF2 expression increased with *KEAP1* knockout.

**Figure 4.**
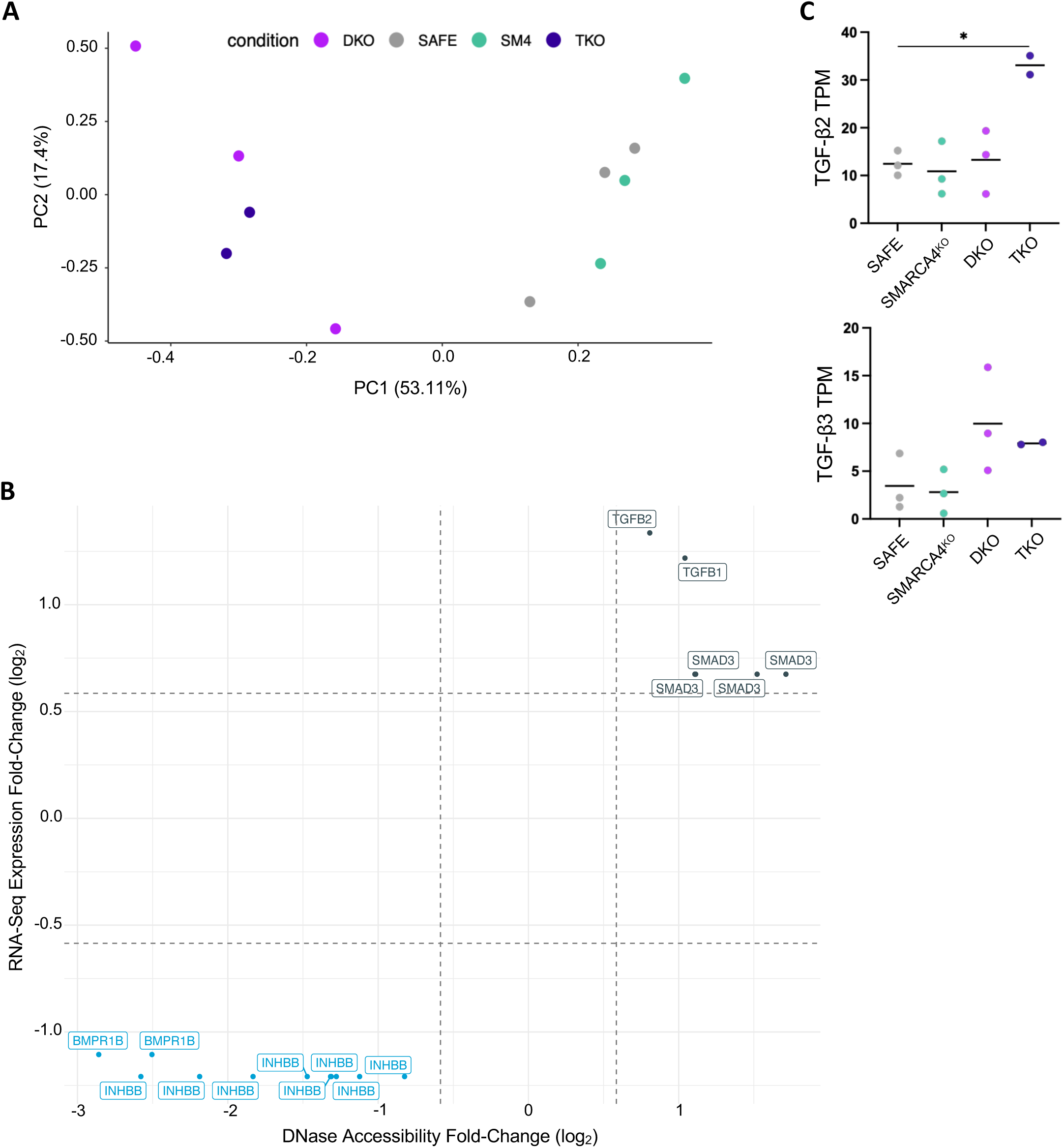
*In vitro* modelling shows *STK11/KEAP1* context-specific TGF-β/EMT pathway expression upon *SMARCA4* loss. **A.** Western blots showing expression of CRISPR/Cas9 knockout (KO) proteins (bolded) and related proteins in H358 single cell clone-derived cell line models. SAFE = sgSAFE KO. Pixel-density of the BRG1 protein band as a percentage of the highest density BRG1 band in the SAFE group (set to 100%) is shown for each lane. Cyclophilin B was probed as a loading control. Gene Set Enrichment Analysis (GSEA) of differentially expressed genes (DEGs) between **B.** SMARCA4^KO^ vs. SAFE and **C.** TKO vs. DKO H358 models. All pathways shown are significantly enriched (adjusted p<0.05) and from the KEGG, GO, and mSigDB gene set databases. NES = normalized enrichment score. See also Additional Files 4 and 5. **D.** Heat map of EMT transcription factors (EMT-TFs) from isogenic clone model RNA-Seq. Hierarchical clustering (supervised by KO genotype group) was performed using Manhattan distance and the Ward.D clustering method. Shading corresponds to Z-score of gene transcripts per million (TPM). **E.** Western blot of expression of TGF-β signaling effectors, EMT-TFs, and associated markers in isogenic clone models. **F.** Bar plot showing concentrations of secreted TGF-β1 ligand as captured by ELISA. Concentrations are averages of 3 technical replicates. Data are represented as mean (± SEM) and significance was calculated by a two-sample unpaired T-test with Bonferroni correction for 6 pairwise comparisons; *p<0.008, **p<0.002. **G.** Dot plot of Transcripts Per Million (TPM) for TGF-β1 (*TGFB1*) by RNA-Seq. Each dot is a model clone. Data are represented as mean (± SEM) and significance was calculated as in Figure 4F. *p<0.008.

Next, we performed RNA-Seq in the isogenic models for transcriptomic analysis. Principal Components Analysis (PCA) of global gene expression data revealed that most variance was imparted by the double *STK11/KEAP1* knockout **(Supplementary Figure S4A)**. DGE and pathway enrichment analyses in the SMARCA4^KO^ versus SAFE models revealed upregulation of interferon response genes among the top enriched signatures, consistent with our results in the clinical RNA-Seq **(Figure 4B; Additional File 4, Tables S1-4)**. Other observations made in the clinical samples were not recapitulated in the SMARCA4^KO^ and SAFE models, including upregulation of OXPHOS. This may reflect the distinct oxidative stress of *in vitro* cell culture versus the relatively hypoxic tumor microenvironment *in vivo*.

In the TKO versus DKO model comparison, we also observed shared and divergent patterns of gene expression with our clinical RNA-Seq **(Figure 4C; Additional File 5, Tables S1-4)**. Interferon signaling was upregulated in the clinical samples but downregulated in the *in vitro* models, while upregulation of cell cycling pathways was shared by both datasets. Notably, upregulation of TGF-β- and EMT-associated gene sets emerged as top enriched signatures in the TKO models, echoing the clinical RNA-Seq and consistent with the known role of the TGF-β pathway as promoter of EMT and metastasis^72,73^. As BRG1-containing SWI/SNF complexes are known to influence the transcriptional repertoire of TGF-β^74–76^, we additionally analyzed the TKO and DKO models by DNaseI-Seq and identified overlapping DNA accessibility and gene expression trends in TGF-β pathway genes **(Supplementary Figure S4B; Additional File 6, Table S1)**. Altogether, these data suggest that the TKO models harbor a unique transcriptional profile, characterized by upregulated TGF-β signaling effectors and TGF-β-modulated EMT transcription factors (EMT-TFs), validating this finding in the clinical RNA-Seq.

TGF-β pathway activity initiates with paracrine secretion of TGF-β ligands, whose binding to cognate TGF-β receptors activates downstream receptor-regulated SMAD proteins (R-SMADs) and ultimately modulates gene expression^72,77^. Protein-level assessment of intracellular signaling effectors downstream of TGF-β receptor (TGFBR) activation revealed similar total expression of SMAD2, SMAD3, and SMAD4 across all model clones **(Figure 4E)** but increased SMAD2 phosphorylation in the TKO clones. Additionally, we observed increased expression of TGF-β ligands TGF-β1/2/3 in the TKO clones, suggesting paracrine ligand secretion as a driver of heightened pathway activity.

We also probed for EMT-TFs SNAI1 and ZEB1, members of the SNAIL and ZEB transcription factor families that are directly upregulated by SMAD-mediated transcription, and EMT markers N-cadherin, E-cadherin, and vimentin^73,78^. Expression of SNAI1 was higher in the DKO clones than in the SMARCA4^KO^ and SAFE clones, and increased further in the TKO clones, which also strongly upregulated ZEB1 **(Figure 4E)**. This could indicate that *STK11* and *KEAP1* loss trigger an initial shift towards a mesenchymal phenotype that is further realized with *SMARCA4* loss—such intermediate “hybrid EMT” cell states have been observed in *in vitro* and *in vivo* NSCLC models^79–81^. The highest expression of EMT markers was seen in the TKO clones, including N-cadherin and vimentin, alongside reduced expression of E-cadherin.

We next performed an ELISA to assess TGF-β1 ligand secretion across our models. Both DKO and in particular TKO cells secreted significantly elevated TGF-β1 relative to SAFE cells **(Figure 4F)**. As the ELISA only measured levels of secreted TGF-β1, we examined the transcript levels of all three TGF-β ligands (TGF-β1, TGF-β2, and TGF-β3) across the models **(Figure 4G; Supplementary Figure S4C)**, observing that TGF-β1 and TGF-β2 transcripts were also highest in the TKO clones. These data suggest that the H358 TKO clones might increase signaling through the TGF-β pathway by secreting higher quantities of TGF-β ligands.

Taken together, multi-omic profiling in clinical samples and isogenic clone model analyses nominated TGF-β pathway upregulation, increased TGF-β ligand secretion, and a pro-metastatic phenotype as characteristic features of *SMARCA4/STK11/KEAP1* triple mutant LUAD.

### H358 TKO models display EMT-associated features and enhanced metastatic capacity

To capture additional *SMARCA4*-specific differences across the isogenic models, we performed a series of phenotypic assays. We observed that H358 TKO clones in culture had a distinctly elongated and flattened appearance **(Figure 5A),** previously observed in LUAD cells with BRG1 knockdown^39^. There were no significant differences in *in vitro* doubling rate across the H358 models **(Supplementary Figure S5A)**. However, we observed that while the SAFE and SMARCA4^KO^ clones did not demonstrate substantial migration or invasion *in vitro*, the DKO clones had greater migration and invasion (p<0.001 and p<0.008 respectively), and the TKO clones displayed a high capacity for migration and invasion (compared to DKO, p<0.0002 and p<0.02 respectively) **(Figure 5B)**.

**Figure 5.**
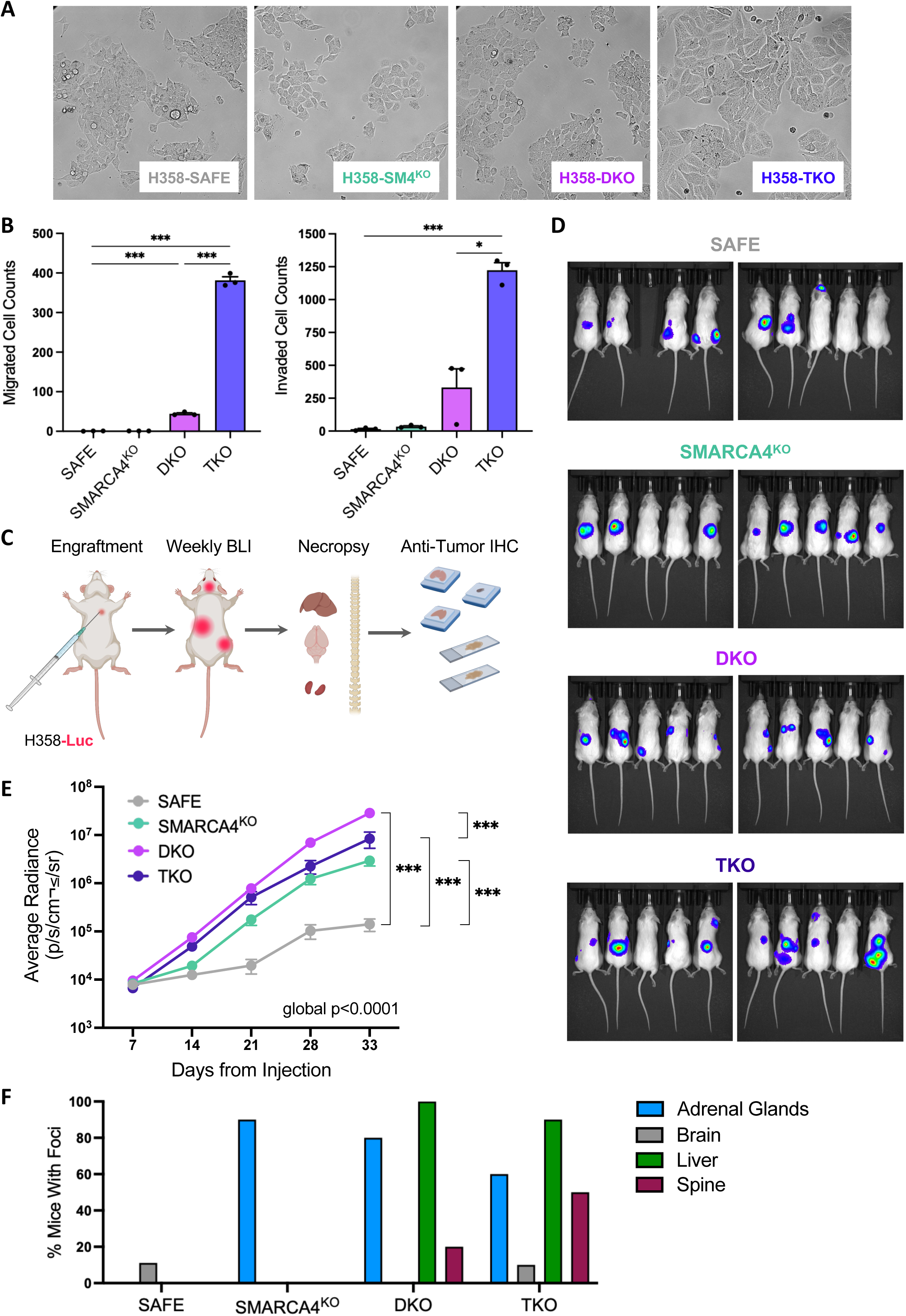
TKO models demonstrate distinct EMT-associated and metastatic phenotypes *in vitro* and *in vivo.* **A.** Bright-field microscope images showing morphology of representative model clones (one per KO genotype group). **B.** Bar plots of migration (left) and invasion (right) assays. Cell numbers are summed cell counts in 5 image fields per replicate, with 3 technical replicates per KO genotype. Data are represented as mean ± SEM. Significance was calculated by a two-sample unpaired T-test with Bonferroni correction for 6 pairwise comparisons; *p<0.008, ***p<0.0002. **C.** Graphical overview of the intracardiac xenograft experiment. **D.** BLI of all mice from the last imaging session prior to sacrifice (Day 33 post-injection). These images are qualitative, not quantitative, depictions of tumor growth. **E.** Growth curves of xenografted tumors per group. Each point is the averaged Average Radiance of all mice per group (n = 9-10) at the indicated time point post-injection. Data are represented as mean ± SEM. Significance was calculated by a general least squares (GLS) approach with Bonferroni correction for 6 pairwise comparisons; ***p<0.0002. **F.** Bar plot of the percentage of mice per model group with at least one tumor in the indicated sites.

Next, we xenografted luciferase-expressing variants of our isogenic cell line models to immunodeficient mice via intracardiac injection, collecting organs representing common LUAD metastatic sites at day 33 to assess metastatic colonization **(Figure 5C)**. We monitored metastatic spread and total tumor growth over the course of the experiment by bioluminescence imaging (BLI) **(Figure 5D)**. SMARCA4^KO^, DKO, and TKO tumors all grew significantly faster than isogenic SAFE controls, as measured by luminescence (p<0.002 for all) **(Figure 5E)**. While the SMARCA4^KO^, DKO, and TKO models did not demonstrate a strong growth advantage *in vitro*, these results suggest that the knockouts may support more rapid metastatic colonization.

To quantitatively assess metastasis, we immunostained the formalin-fixed, paraffin-embedded organ tissue with an anti-human mitochondrial antigen antibody to visualize model-derived tumors and tallied the number of organs with at least one tumor. We then evaluated organotropism by plotting the incidence of metastasis in the sites we surveyed for each model. Metastases were detected for each model with the knockout models displaying the highest frequencies of metastasis, as captured by BLI **(Figure 5F)**. TKO model mice had the greatest breadth of metastatic sites, followed by the DKO model mice, with only one metastatic site each captured in SAFE and SMARCA4^KO^ model mice. These observations suggest that H358 cells display broader organotropism (metastasize to a wider variety of sites) with the addition of the DKO and TKO, aligning with our observation of higher metastatic burden in triple mutant patients **(Figure 2C; Supplementary Figure S2B)**.

## DISCUSSION

Our clinicogenomic analysis captured previously reported co-occurrence of *SMARCA4* loss, *STK11/KEAP1* loss, and loss of all three genes with canonical LUAD genetic drivers and identified new associations, particularly in triple mutant tumors. We additionally observed that *SMARCA4/STK11/KEAP1* co-loss correlated with poor prognosis that might be driven by increased metastatic burden, and that these patients appear to have poor outcomes when treated with either chemotherapy or immunotherapy. Through gene-level analysis, we confirmed co-losses of heterozygosity (co-LOH) in these three genes as a frequent event in LUAD and captured enrichment for biallelic gene loss of *SMARCA4* in triple mutant tumors. *SMARCA4* mutations that are predicted to have most functionally disruptive effects (e.g. deletions, truncations) or to result in biallelic *SMARCA4* loss have been correlated to the worst outcomes in patients with *SMARCA4*-deficient LUAD^8,82^. In addition, *STK11* and *KEAP1* mutations have been observed to occur significantly more frequently in *SMARCA4*-mutant tumors than wild-type tumors, and even more frequently in *SMARCA4*-mutant tumors with severely disruptive *SMARCA4* mutations^8^. It is possible that a higher preponderance of highly disruptive *SMARCA4* mutations in triple mutant LUAD patients contributes to their exceptionally poor prognosis.

Given the high frequency of co-LOH of *SMARCA4, STK11*, and *KEAP1* in LUAD, it is probable that studies correlating their losses with clinical outcomes would draw different interpretations if triple mutant patients are not considered separately. For example, sequencing studies re-examining the reported correlation between *STK11* and *KEAP1* mutations and worse immunotherapy response in LUAD have found this correlation to be stronger when these mutations are accompanied by oncogenic KRAS^25,29^ or low TMB^83^. Our findings suggest that conflicting findings on the correlation of *SMARCA4* deficiency with chemotherapy response may stem from the absence of *SMARCA4, STK11*, and *KEAP1*-status defined comparator groups in these studies^40,84^, likely due to the challenge of assembling sufficient patient numbers in each group to enable statistical comparisons. A recent immunotherapy study which accounted for these co-incident mutations reinforces this notion, as the authors found that, when analyzed separately from *STK11/KEAP1*-mutant and *SMARCA4/STK11/KEAP1*-mutant patients, *SMARCA4* only mutant patients had significantly longer progression-free survival and numerically longer overall survival on immune checkpoint inhibitor (ICI) therapy^42^, contradicting previous studies that examined *SMARCA4*-mutant tumors without respect to *STK11/KEAP1* co-mutant status^8,41,43^. Indeed in our own study, we found that *SMARCA4* only mutant LUAD patients fared similarly on ICI to patients who were wild-type for all three genes. Our data add to a growing body of evidence that the implications of *SMARCA4* deficiency in LUAD are highly context-dependent and particularly impacted by co-mutation of *STK11* and *KEAP1*. LUAD sequencing studies with stratification by *STK11* and *KEAP1* status have enabled characterization of these genes as predictive biomarkers and the recent identification of dual immune checkpoint blockade as a potentially effective therapeutic approach^85^. We anticipate that the increasing accessibility of multi-institutional sequencing datasets (e.g. AACR Project GENIE) will potentiate investigation of *SMARCA4*-deficient LUAD while accounting for *STK11* and *KEAP1* co-loss, advancing both our clinical understanding of triple mutant LUAD and our efforts to provide precision medicine to a wider fraction of LUAD patients.

Our transcriptomic analyses of LUAD clinical samples indicated that *SMARCA4*-mutant tumors, independent of their *STK11/KEAP1* background, show upregulation of interferon (IFN)-α and -γ pathway genes and downregulation of alveolar type II (AT2) cell lineage markers. *SMARCA4*-deficient lung cancers are highly undifferentiated by histological evaluation^69,70^, and we observed the greatest loss of these markers in triple mutant tumors. Our observation of upregulated TGF-β gene sets in the same tumors suggests that TGF-β activity may have a role in this phenotype, as EMT induction can trigger cancer stem cell formation and impart stem cell-like features^79^.

Given the inherent heterogeneity of clinical cohorts, we sought to assess which observations from our clinical datasets could be validated by comparisons to isogenic models, which we generated from the *KRAS^G12C^*-mutant LUAD cell line NCI-H358. Bulk RNA-Seq of these models revealed upregulation of TGF-β gene expression as a shared feature of our triple mutant clinical samples and triple knockout models, suggesting that this finding might be specific to co-loss of *SMARCA4/STK11/KEAP1*. RAS/MAPK pathway signaling is required for TGF-β-mediated induction of EMT^77,86–90^, and the acquisition of oncogenic KRAS by epithelial cells can accelerate their malignant transformation and impart metastatic phenotypes through TGF-β-stimulated EMT^88,90–93^. The presence of oncogenic *KRAS* might prime H358 cells to undergo EMT following upregulation of TGF-β and/or altered transcriptional activity through SMADs in association with BRG1 loss. However, the presence of EMT transcriptional signatures in the clinical RNA-Seq, which was driver-agnostic, suggests that this effect might not be limited to *KRAS*-mutant LUAD. Further investigation will clarify whether upregulation of EMT genes is a convergent outcome of triple mutant LUAD biology, regardless of driver status, or whether RAS activity is otherwise increased to enact TGF-β-induced EMT in in *KRAS* wild-type triple mutant LUAD.

We additionally captured *SMARCA4*-specific overlapping chromatin accessibility and gene expression changes in our triple knockout models. BRG1-containing SWI/SNF complexes directly modulate TGF-β-mediated transcription by cooperatively binding with SMAD complexes and histone acetyltransferases (e.g. CBP, p300) and relaxing chromatin at SMAD-regulated promoters^74,75^. In keeping with this, loss of BRG1 has been observed to alter selectivity of SMADs for a subset of TGF-β target genes^76^. Whether the unique activation of TGF-β signaling and gene expression observed in our triple knockout models arises from altered SMAD targeting warrants further investigation.

We observed that *SMARCA4* knockout induces TGF-β signaling and EMT marker expression most strongly in the context of concomitant loss of *STK11* and *KEAP1*. It is possible that elevated TGF-β activity is one of many paths to EMT for *SMARCA4*-deficient LUAD cells. Concepción, et al. found that homozygous deletion of Smarca4 in KP model primary tumors produced loss of lung lineage factors with increased expression of genes related to metastasis and high grade malignancy. Upon metastasis, these clones assumed diverse transcriptional programs in transit to their eventual metastatic sites, suggesting that *SMARCA4* loss lifts the epigenetic constraints of lung cell lineage and enhances capacity for transcriptional reprogramming^37^. This plasticity may facilitate or expedite adoption of an EMT transcriptional program, which can be stimulated through various paracrine signaling inputs^79^. In this scenario, *STK11* and *KEAP1* inactivation may constrict the array of transcriptional outcomes potentiated by *SMARCA4* loss, shunting these cells towards TGF-β-driven EMT.

Echoing our clinicogenomic results, triple knockout of *SMARCA4, STK11*, and *KEAP1* appeared to enhance metastatic capacity *in vitro* and *in vivo*, in keeping with the elevated TGF-β signaling and EMT marker expression captured in these models. TGF-β-induced EMT supports intravasation and dissemination of cells from primary tumors by disrupting cell adhesion and polarity and imparting mesenchymal-like migratory and invasive phenotypes^73,79^. Future studies will be crucial to evaluating TGF-β inhibition as a potential therapeutic strategy in *SMARCA4/STK11/KEAP1* co-mutant LUAD, in combinations with platinum-based chemotherapy, targeted therapy (e.g. KRAS^G12C^ inhibition), or immunotherapy.

TGF-β activity in *SMARCA4/STK11/KEAP1* co-mutant LUAD might also contribute to poor outcome through adverse effects on immunosurveillance. TGF-β signaling has been shown to facilitate immunoevasion of circulating tumor cells from surveilling NK cells, promote quiescence and reduced expression of MHC Class I and NK receptor ligands by disseminated clones, and maintain an immunosuppressive, T-cell-depleted tissue microenvironment surrounding metastatic colonies upon their outgrowth^72,73,94,95^. In a mouse model of LUAD, TGF-β secretion by host tissue was observed identified to suppress STING signaling and maintain dormancy of metastatic cells, and STING agonist treatment restored their cell cycling and enabled their killing by T-cells and NK cells^96^. To fully understand the consequences of TGF-β signaling on development and progression of triple mutant tumors, and to assess the potential efficacy of combining TGF-β inhibition and immunotherapy in this context, investigation in immunocompetent models is essential.

## CONCLUSIONS

In this study, we synthesized clinicogenomic data, transcriptional profiling, and *in vitro* and *in vivo* phenotypic assays to characterize *SMARCA4* loss in LUAD, both on its own and in the clinically-informed context of co-occurring *STK11* and *KEAP1* inactivation. We identified shorter overall survival, increased metastatic burden and spread, enhanced metastasis-associated phenotypes, and upregulation of TGF-β and EMT transcriptional signatures as distinguishing features of combined *SMARCA4/STK11/KEAP1* loss in LUAD. Given the lack of targeted therapy and poor prognosis for patients with *SMARCA4/STK11/KEAP1* triple mutant LUAD, these findings establish a rationale to investigate the mechanistic origins of TGF-β upregulation and dependence in this setting, clarify the link between TGF-β activity and metastasis-associated phenotypes, and consider triple mutant patients as a clinically distinct and prognostically meaningful subset of LUAD in future clinicogenomic studies.

## ABBREVIATIONS

AT2: alveolar type II
ATM: ataxia-telangiectasia mutated
ATP: adenosine triphosphate
BLI: bioluminescence imaging
BRG1: Brahma-related gene 1
BRM: Brahma
CNS: central nervous system
CRISPR/Cas9: Clustered regularly interspaced palindromic repeats/CRISPR associated protein 9
DGE: differential gene expression
DKO: double knockout
DNA: deoxyribonucleic acid
EGFR: epidermal growth factor receptor
ELISA: enzyme-linked immunosorbent assay
EMT: epithelial-mesenchymal transition
FACETS: Fraction and Allele-Specific Copy Number Estimates from Tumor Sequencing
GEMM: genetically-engineered mouse model
GSEA: gene set enrichment analysis
ICI: immune checkpoint inhibitor
IFN: interferon
IO: immunotherapy
KEAP1: Kelch-like ECH-associated protein 1
KO: knockout
KP: Kras^LSL(lox–stop–lox)-G12D/+^ Trp53^fl/fl^
KRAS: Kirsten rat sarcoma
LKB1: liver kinase B1
LN: lymph node
LOH: loss of heterozygosity
LUAD: lung adenocarcinoma
MAPK: mitogen-activated protein kinase
MOI: multiplicity of infection
MSK-IMPACT: Memorial Sloan Kettering-Integrated Molecular Profiling of Actionable Cancer Targets
NES: normalized enrichment score
NRF2: nuclear factor erythroid 2-related factor 2
NSCLC: non-small cell lung cancer
NSG: NOD.Cg-Prkdcscid Il2rgtm1Wjl/SzJ
OR: odds ratio
ORR: overall response rate
OS: overall survival
OXPHOS: oxidative phosphorylation
PCA: Principal Components Analysis
QC: quality control
pRB: retinoblastoma protein
R-SMAD: receptor-regulated SMAD
RNA: ribonucleic acid
ROI: region of interest
sgRNA: small guide RNA
sgSAFE: safe-targeting guide RNA
SIK: salt-inducible kinase
SMAD: mothers against decapentaplegic homolog
SMARCA2: SWI/SNF related, matrix associated, actin dependent regulator of chromatin, subfamily a, member 2
SMARCA4: SWI/SNF related, matrix associated, actin dependent regulator of chromatin, subfamily a, member 4
SNAI1: snail family transcriptional repressor 1
STK11: serine/threonine kinase 11
STING: stimulator of interferon genes
SWI/SNF: SWItch/Sucrose Non-Fermentable
TCGA: The Cancer Genome Atlas
TF: transcription factor
TGF-β/TGFB: transforming growth factor beta
TGFBR: TGF-β receptor
TIDE: Tracking of Indels by Decomposition
TKO: triple knockout
TMB: tumor mutational burden
TP53: tumor protein P53
TPM: transcripts per million
WT: wildtype
ZEB1: zinc finger E-box-binding homeobox 1

## DECLARATIONS

### Ethics Approval

This study was approved by the Memorial Sloan Kettering Cancer Center (MSKCC) Institutional Review Board under protocols 12-245, 14-209, and 16-1566. All patients included in the study had tumors analyzed prospectively by MSK-IMPACT as part of routine clinical care at MSKCC. The detailed review of demographic, radiologic and clinical information was performed retrospectively. All animal experiments were approved by the Memorial Sloan Kettering Cancer Center (MSKCC) Animal Care and Use Committee.

### Consent for Publication

Not applicable.

### Availability of Data and Materials

The *in vitro* RNA-Seq and DNAseI-Seq and clinical RNA-Seq datasets generated and analyzed during the current study are publicly available in the gene expression omnibus (GEO) under accession number GSE299604.

### Authors Contributions

Conceptualization: EC, CMR, AQV. Methodology: EC, CW, BPM, ER, HS, DK, ES, WK, NF, AF, MCM, IL, UB, KL, DB, AE; Investigation: EC, BPM, AQV, CMR; Validation: EC, AQV, CMR; Formal Analysis: EC, SN, ST, YAZ, JJ, JL, WC, AA; Writing – Original Draft: EC; Review & Editing: All authors; Supervision: AQV and CMR; Funding acquisition: CMR. All authors read and approved the final version of the manuscript.

## Acknowledgements

The authors would like to thank many cores at Memorial Sloan Kettering for their contributions to key experiments in this manuscript, namely: Maria Corazón Mariano of the Precision Pathology Biobanking Center; Kevin Chen, Phoebe Bacchus, and Surahbi Parte of the Antitumor Assessment Core; Eric Chan and Eric Rosiek of the Molecular Cytology Core; Sébastien Monette, Sebastian Carrasco, and Mohamed Atmane of the Laboratory of Comparative Pathology; and the Integrated Genomics Operation Core. We also thank Christina Wilson and Natasha Rekhtman of the Thoracic Pathology Service for their assistance in coordinating clinical aspects of the project. We thank JT Poirier and Yuan Hao for data analysis advice and all members of the Rudin Lab for their support, both technical and personal. For MSK-IMPACT and the Molecular Cytology Core, we acknowledge the Center Core grant (P30 CA008748). For MSK-IMPACT, we also acknowledge the Molecular Diagnostics Service in the Department of Pathology, and the Marie-Josee and Henry R. Kravis Center for Molecular Oncology. We acknowledge Genewiz for performing next-generation RNA sequencing and Sanger sequencing and Biorender for use of their illustrations in figures.

## AUTHOR INFORMATION

Álvaro Quintanal-Villalonga and Charles M. Rudin have a shared senior authorship.

## ADDITIONAL FILES

**Additional File 1.** Supplementary methods Tables S1-8

**Additional File 2.** SMARCA4 Only Mutant vs. Unaltered Clinical GSEA Tables S1-4

**Additional File 3.** Triple Mutant vs. Double Mutant Clinical GSEA Tables S1-4

**Additional File 4.** SMARCA4^KO^ vs. SAFE H358 Model GSEA Tables S1-4

**Additional File 5.** TKO vs. DKO H358 Model GSEA Tables S1-4

**Additional File 6.** TKO vs. DKO H358 Model RNA-Seq and DNaseI-Seq Integration Table S1

