## Supplementary Figures and Methods for "*SMARCA4, STK11,* and *KEAP1* co-inactivation associates with poor prognosis and upregulation of the TGF-β pathway in lung adenocarcinoma"

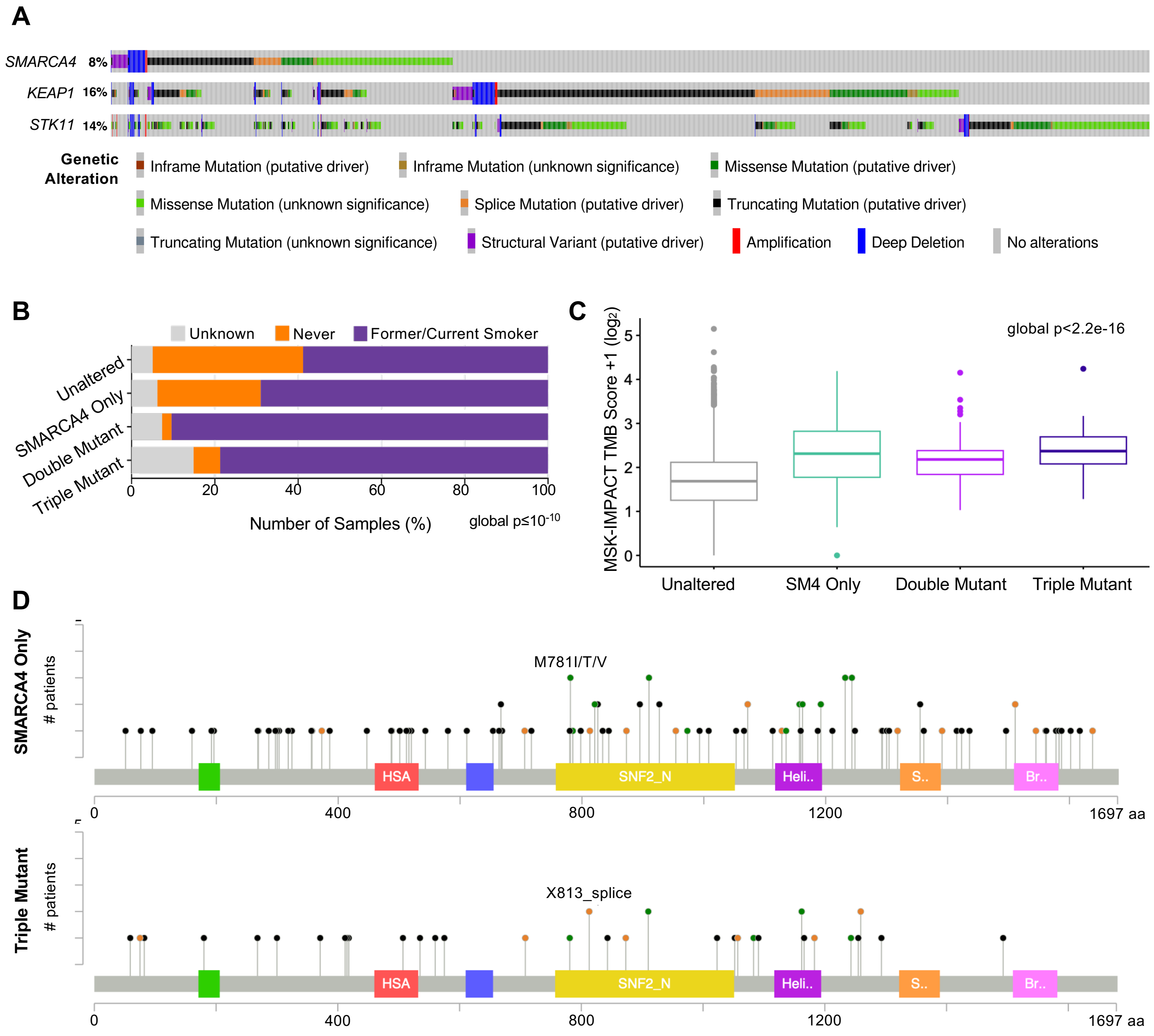
A
SMARCA4 8%
KEAP1 16%
STK11 14%
Genetic Alteration
Inframe Mutation (putative driver)
Inframe Mutation (unknown significance)
Missense Mutation (putative driver)
Missense Mutation (unknown significance)
Splice Mutation (putative driver)
Truncating Mutation (putative driver)
Truncating Mutation (unknown significance)
Structural Variant (putative driver)
Amplification
Deep Deletion
No alterations
B
Unknown
Never
Former/Current Smoker
Unaltered
SMARCA4 Only
Double Mutant
Triple Mutant
0
20
40
60
80
100
Number of Samples (%)
global p≤10-10
C
global p<2.2e-16
MSK-IMPACT TMB Score +1 (log2)
0
1
2
3
4
5
Unaltered
SM4 Only
Double Mutant
Triple Mutant
D
SMARCA4 Only
### patients
M781I/T/V
HSA
SNF2_N
Heli..
S..
Br..
0
400
800
1200
1697 aa
Triple Mutant
### patients
X813_splice
HSA
SNF2_N
Heli..
S..
Br..
0
400
800
1200
1697 aa

**Figure S1. Related to Figure 1. MSK-IMPACT cohort clinical characteristics, co-altered genes, and *SMARCA4* mutational spectrum. A.** MSK-IMPACT OncoPrint of *SMARCA4*, *STK11*, and *KEAP1* genetic alterations from tumor sequencing of 9,374 lung adenocarcinoma patients. Each vertical line is a single patient. Tumors that were wild-type for all three genes are not shown. **B.** Stacked bar plot showing the distribution of various smoking statuses (Former/Current Smoker, Never or Unknown) across patients in the clinicogenomic cohort. The p-value was calculated by chi-squared test. **C.** Box plot depicting tumor mutational burden (TMB) Scores in samples from each cohort group. The center line is the median score. The p-value was calculated by Wilcoxon test. SM4 = SMARCA4. **D.** Lollipop plot of the *SMARCA4* gene depicting loci of hotspot mutations in samples from the SMARCA4 Only and Triple Mutant groups. Annotated domains from left to right are: QLQ (green), HSA (red), BRK (blue), SNF2 N-terminal (yellow), helicase (purple), SnAC (orange) and bromodomain (pink). Lollipop colors correspond to the following mutation types: black = truncating, green = missense, orange = splice. aa = amino acids.

**A**

| Group | Number at Risk | | | | |
|---|---|---|---|---|---|
| Unaltered(n=4618) | 4618 | 2039 | 800 | 125 | 0 |
| SMARCA4 Only(n=129) | 129 | 42 | 16 | 5 | 0 |
| Double Mutant(n=175) | 175 | 60 | 25 | 7 | 0 |
| Triple Mutant(n=47) | 47 | 5 | 1 | 0 | 0 |
| Months | 0 | 30 | 60 | 90 | 120 |

**B**

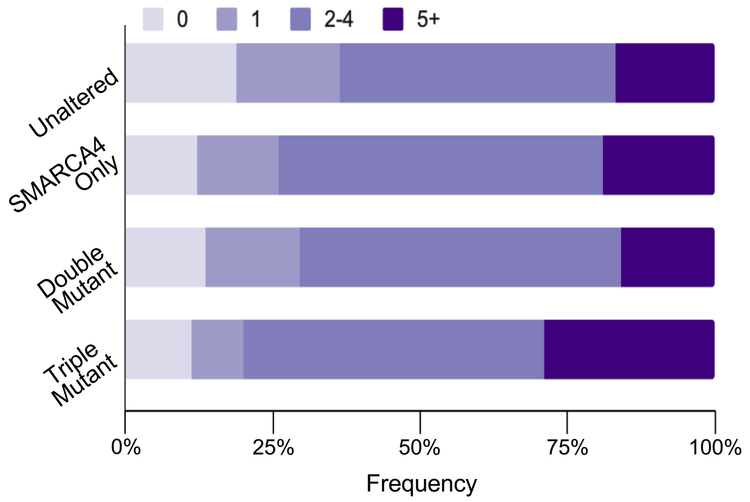


**C**

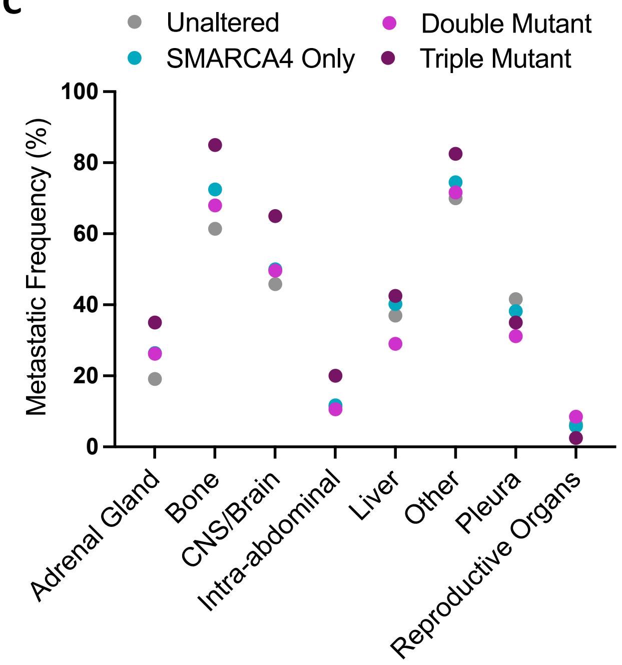


**D**

| Group (Chemo cohort) | Number at Risk | | | |
|---|---|---|---|---|
| Unaltered(n=2821) | 2821 | 1064 | 655 | 466 |
| SMARCA4 Only(n=104) | 104 | 33 | 15 | 10 |
| Double Mutant(n=81) | 81 | 20 | 11 | 8 |
| Triple Mutant(n=23) | 23 | 3 | 1 | 0 |
| Months | 0 | 10 | 20 | 30 |

**E**

| Group (IO cohort) | Number at Risk | | | |
|---|---|---|---|---|
| Unaltered(n=1795) | 1795 | 763 | 428 | 271 |
| SMARCA4 Only(n=76) | 76 | 39 | 24 | 19 |
| Double Mutant(n=79) | 79 | 24 | 16 | 9 |
| Triple Mutant(n=16) | 16 | 1 | 0 | 0 |
| Months | 0 | 10 | 20 | 30 |

**Figure S2. Related to Figure 2. Per patient metastasis site counts across clinicogenomic cohort groups. A.** Patient numbers for the overall survival Kaplan-Meier curves in Figure 2A. **B.** Stacked bar plots depicting how frequently patients in each cohort group have 0, 1, 2-4, or 5+ reported sites of metastasis. **C.** Dot plot showing the frequencies of patients per cohort group with metastases at the listed metastatic sites. Secondary lung metastases and lymph node metastases were excluded from this analysis. Patient numbers for the corresponding survival Kaplan-Meier curves in **D.** Figure 2D and **E.** Figure 2E.

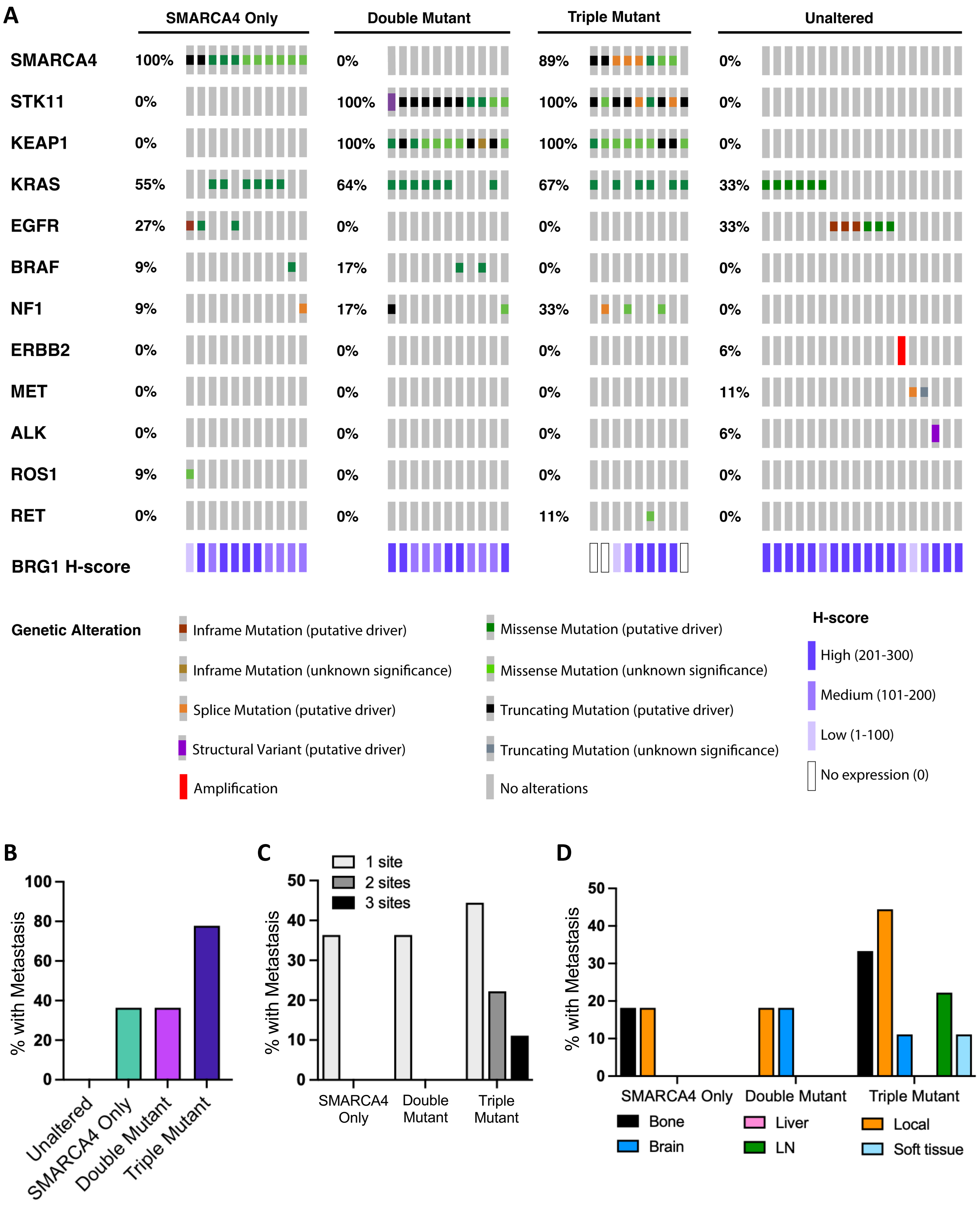
A
SMARCA4 Only
Double Mutant
Triple Mutant
Unaltered
SMARCA4 100% 0% 89% 0%
STK11 0% 100% 100% 0%
KEAP1 0% 100% 100% 0%
KRAS 55% 64% 67% 33%
EGFR 27% 0% 0% 33%
BRAF 9% 17% 0% 0%
NF1 9% 17% 33% 0%
ERBB2 0% 0% 0% 6%
MET 0% 0% 0% 11%
ALK 0% 0% 0% 6%
ROS1 9% 0% 0% 0%
RET 0% 0% 11% 0%
BRG1 H-score
Genetic Alteration
Inframe Mutation (putative driver)
Inframe Mutation (unknown significance)
Splice Mutation (putative driver)
Structural Variant (putative driver)
Amplification
Missense Mutation (putative driver)
Missense Mutation (unknown significance)
Truncating Mutation (putative driver)
Truncating Mutation (unknown significance)
No alterations
H-score
High (201-300)
Medium (101-200)
Low (1-100)
No expression (0)
B
% with Metastasis
100
80
60
40
20
0
Unaltered
SMARCA4 Only
Double Mutant
Triple Mutant
C
1 site
2 sites
3 sites
% with Metastasis
50
40
30
20
10
0
SMARCA4 Only
Double Mutant
Triple Mutant
D
% with Metastasis
50
40
30
20
10
0
SMARCA4 Only
Double Mutant
Triple Mutant
Bone
Brain
Liver
LN
Local
Soft tissue

**Figure S3. Related to Figure 3. Genetic and clinical overview of the clinical RNA-Seq cohort. A.** MSK-IMPACT OncoPrint of genetic alterations in the clinical sample cohort. Each column is a single sample, and there is no sample overlap among the four cohort groups (SMARCA4 Only, Double Mutant, Triple Mutant, Unaltered). Purple-shaded rectangles along the bottom reflect the BRG1 H-score of each sample as assessed from BRG1 immunostaining of tumor-matched FFPE samples. Scores are categorized as high (H-score = 201-300), medium (101-200), and low (1-100), with an H-score of 0 indicating no BRG1 staining. **B.** Bar plot showing the percentage of patients with recorded metastasis in each group. **C.** Bar plot showing the percentages of patients with 1, 2, or 3 metastatic sites. **D.** Bar plot of percentages of patients per group with at least one recorded met in the listed sites: bone, brain, liver, lymph nodes (LN), local (lung), and soft tissue.

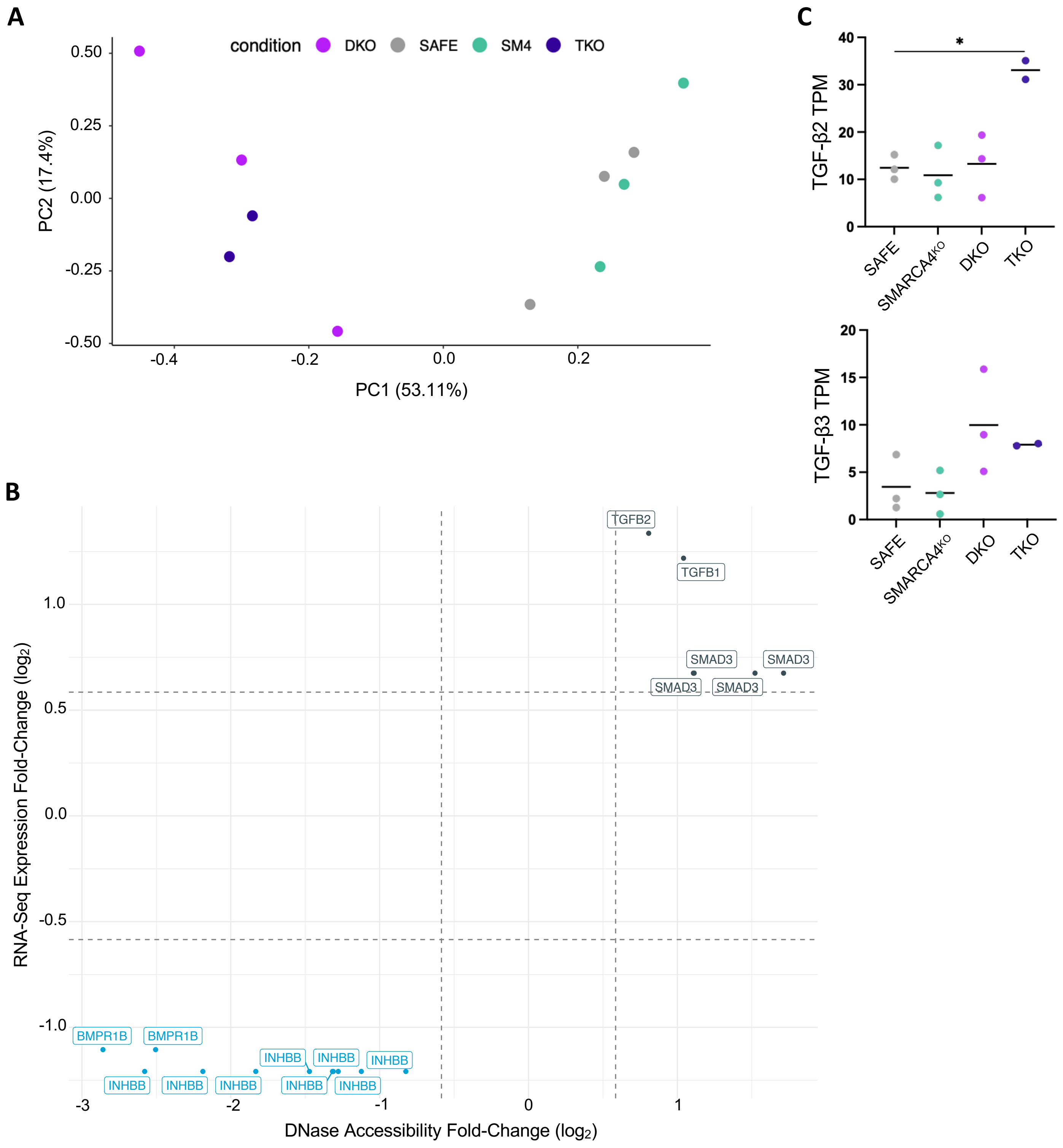
A
condition DKO SAFE SM4 TKO
PC2 (17.4%)
0.50
0.25
0.00
-0.25
-0.50
-0.4
-0.2
0.0
0.2
PC1 (53.11%)
B
RNA-Seq Expression Fold-Change (log2)
1.0
0.5
0.0
-0.5
-1.0
TGFB2
TGFB1
SMAD3
SMAD3
SMAD3
SMAD3
BMPR1B
BMPR1B
INHBB
INHBB
INHBB
INHBB
INHBB
INHBB
INHBB
INHBB
-3
-2
-1
0
1
DNase Accessibility Fold-Change (log2)
C
TGF-β2 TPM
40
30
20
10
0
*
SAFE
SMARCA4KO
DKO
TKO
TGF-β3 TPM
20
15
10
5
0
SAFE
SMARCA4KO
DKO
TKO

**Figure S4. Related to Figure 4. *In vitro* H358 model RNA-Seq PCA and DNase-Seq. A.** Principal Component Analysis (PCA) of isogenic clone models in RNA-Seq. Variance percentages are based on gene transcripts per million (TPMs). SM4 = SMARCA4 $KO^{KO}$. **B.** Plotted RNA-Seq gene expression and DNaseI-Seq gene accessibility fold-changes for genes in the KEGG "TGF-beta Signaling Pathway" gene set between the TKO versus DKO H358 models. Genes shown have fold-changes in accessibility and gene expression $\geq \log_2(1.5)$ and are significantly differentially expressed and accessible (adjusted p<0.5) by RNA-Seq and DNaseI-Seq. See also Additional File 6. **C.** Dot plots of Transcripts Per Million (TPM) counts for TGF-β2 (*TGFB2*) and TGF-β3 (*TGFB3*) by RNA-Seq. Each dot is a single cell clone. Data are represented as mean (± SEM) and significance was calculated by a two-sample unpaired T-test with Bonferroni correction for 6 pairwise comparisons; *p<0.008.

A

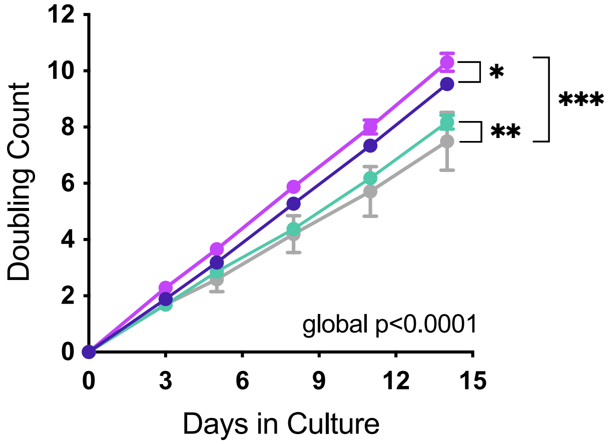

Doubling Count
12
10
8
6
4
2
0
0
3
6
9
12
15
Days in Culture
global p<0.0001
*
**
***

**Figure S5. Related to Figure 5. H358 model *in vitro* and *in vivo* growth and metastasis phenotypes. A.** Growth curves of H358 single cell clone models over two weeks in *in vitro* culture. At each time point, cells were counted and reseeded until the end of the assay to calculate doublings. Model clones were biological replicates, and curves depict the mean growth of clones (measured in cell doublings) with the indicated model genotype across the assay. Data are mean ± SEM. Significance was calculated by a general least squares (GLS) approach with Bonferroni correction for 6 pairwise comparisons; *p<0.008, **p<0.002, ***p<0.0002. ns = not significant.

#### SUPPLEMENTARY METHODS

##### Smoking Status Analysis

Smoking status (former/current vs. never) was obtained via regular expression extraction applied to the first available clinician assessment for a given patient. The algorithm was created based on a previously published cohort of 247 patients with NSCLC and previously annotated smoking status, withholding data from patients also present in the MSK BioPharma Collective (BPC) NSCLC cohort. The model was validated based on the MSK BPC NSCLC cohort (AACR GENIE). Comparison across groups was done using a chi-squared test.

##### Tumor Mutational Burden Analysis

Tumor mutational burden (TMB) was defined as the count of nonsynonymous mutations per megabase covered by the IMPACT panel. The fraction genome altered was determined by assessing the length of the sequenced genome with a log2 copy number variation (gain or loss) greater than 0.2, divided by the total size of the genome profiled for copy number. Comparison across groups was done using a Wilcoxon test.

##### BRG1 and H&E Immunohistochemistry

To identify potential clinical samples that lacked BRG1 expression and/or had incorrectly annotated histology in MSK-IMPACT, FFPE embedded tumor tissue blocks corresponding to the surgically resected tumor tissues selected for our cohort were immunostained for BRG1 expression. The primary antibody (BRG1 clone G-7, Santa Cruz, catalog no. sc-17796) was used at a 1:400 dilution. Immunohistochemistry was performed on the BOND RX using the bond polymer detection kit (Leica, catalog no. DS9800). The manufacturer's standard protocol was implemented using 30 minutes of heat-induced epitope retrieval with ER2 buffer and 30 minutes of primary antibody incubation. BRG1-stained slides and sample matched hematoxylin and eosin (H&E)-stained slides were evaluated by a pathologist for tumor cell histology and BRG1 expression, and H-scores for BRG1 expression were assigned. Any samples that were not histologically lung adenocarcinoma were removed from the cohort. Samples that were SMARCA4 wild-type but lacked BRG1 expression (H-score=0) were treated as SMARCA4 mutant in the analysis.

**Clinical Sample Preparation**

Upon retrieval, the frozen tissues were OCT embedded, sectioned, and mounted to slides for H&E staining to identify non-necrotic tumor areas and immunostaining with an anti-BRG1 antibody (Santa Cruz, sc-17796). 2-3 3mm punches were collected per sample and stored at -80° C until RNA isolation. For RNA isolation, frozen tissue punches were weighed and 20-30mg were homogenized in RLT. Nucleic acids were extracted using the AllPrep DNA/RNA Mini Kit (Qiagen catalog no. 80204) according to the manufacturer's instructions. RNA was eluted in nuclease-free water and DNA in 0.5X Buffer EB and submitted for bulk RNA-Seq

**Clinical RNA-Seq Library Preparation and Sequencing**

Extracted RNA from frozen tissue samples was submitted for bulk RNA-Seq. Total RNA library preparation and sequencing were conducted at Azenta Life Sciences (South Plainfield, NJ, USA) as follows: RNA samples were quantified using Qubit 2.0 Fluorometer (ThermoFisher Scientific) and RNA integrity was checked with 4200 TapeStation (Agilent Technologies). Samples were initially treated with TURBO DNase (Thermo Fisher Scientific) to remove DNA contaminants. rRNA depletion was performed by using Qiagen FastSelect rRNA HMR Kit (Qiagen). RNA sequencing libraries were prepared using NEBNext Ultra II RNA Library Preparation Kit for Illumina by following the manufacturer's recommendations (NEB, Ipswich). Briefly, enriched RNAs were fragmented for 15 minutes at 94 °C. First strand and second strand cDNA were subsequently synthesized. cDNA fragments were end repaired and adenylated at 3'ends, and universal adapters are ligated to cDNA fragments, followed by index addition and library enrichment with limited cycle PCR.

Sequencing libraries were validated using the Agilent Tapestation 4200 (Agilent Technologies), and quantified using Qubit 2.0 Fluorometer (ThermoFisher Scientific) as well as by quantitative PCR (KAPA Biosystems). The sequencing libraries were multiplexed and clustered onto a flow cell on the Illumina NovaSeq instrument according to manufacturer's instructions. The samples were sequenced using a 2x150bp Paired End (PE) configuration. Image analysis and base calling were conducted by the NovaSeq Control Software (NCS). Raw sequence data (.bcl files) generated from Illumina NovaSeq was converted into fastq files and de-multiplexed using Illumina bcl2fastq 2.20 software. One mismatch was allowed for index sequence identification.

**Lentiviral Production**

Lenti-X™ 293T cells (Takara Bio, catalog no. 632180) were treated with jetPRIME transfection reagent and buffer (Polyplus, catalog no. 114-15) and co-transfected with lentiviral plasmid and packaging vectors (psPAX2 and pMD2.G). The media was changed 16 hours after transfection and collected 72 hours after transfection to recover lentiviruses. Cell debris were removed from the lentiviral supernatant using a 0.45µM syringe filter. Lentiviruses were concentrated by mixing with Lenti-X™ concentrator overnight at 4° C and centrifuging at 1,500 x g for 45 minutes at 4° C. Media was aspirated and lentiviruses were resuspended in 1mL RPMI for storage at -80°C.

**Lentiviral Transduction and Antibiotic Selection**

For each cell line that was transduced, the virus used in transduction was titered in that cell line to ensure transduction at an MOI of 0.3. 1 million cells were seeded to T75 flasks the night before transducing. For transduction, 8µg/mL of polybrene was added to the media (RPMI with 10% FBS and 1X PS) and the lentivirus was thawed at 4° C and kept on ice immediately prior to dispensing onto the cells. Cells were incubated overnight and polybrene-containing media was replaced with fresh media the following day. Antibiotics were added for selection 2 days after transduction, and selection was considered completed when control cells that had received no lentivirus were all observed to be dead under the microscope. The following antibiotic concentrations were used for H358 cells: 2µg/mL of blasticidin, 2µg/mL of puromycin, 200µg/mL of neomycin.

For luciferase (Luc) expressing cell line generation, Individual single cell clone cell lines were transduced with pLL-CMV-rFluc-T2A-GFP-mPGK-Puro (Lenti-Labeler) lentivirus at an MOI of 0.3 and selected with puromycin. Luciferase activity was verified by seeding cells to a white 96-well plate and, the following morning, replacing the media with media containing Bio-Glo™ luciferin (Promega, catalog no. G7941) diluted 1:50. Cells were incubated in luciferase media in the dark for 15 minutes at RT. Luminescence signal was read on a BioTek Synergy Neo microplate reader (Agilent).

**Single Cell Clone Isolation and Screening**

H358 cells transduced with sgSAFE, sgSMARCA4, or sgKEAP1 and sgSTK11 lentiGuide vectors were dilution seeded to 4 96-well plates. The day after seeding, wells containing single cells were marked and incubated until their wells were confluent. Single cell clones were trypsinized and reseeded to a larger well plate, and progressively trypsizined and reseeded to larger plates until they reached confluence in a T25 flask, at which point clones were cryopreserved until screening. Clones were prioritized for screening based on their probability of being a single cell clone (i.e. their well location in the original 96-well plates). *SMARCA4* knockout clones with the most loss of BRG1 expression compared to the parental cell line were selected and Sanger sequenced to verify clonality. sgSAFE clones with the most similar expression of BRG1 as the parental cell line were selected and Sanger sequenced.

**Cell Line RNA-Seq Sample Preparation and Sequencing**

1 million cells were trypsinzed, washed twice with cold PBS, and flash frozen as cell pellets for submission for bulk RNA-Seq. RNA extraction, library preparation, sequencing and analysis was conducted at Azenta Life Sciences (South Plainfield, NJ, USA) as follows: Total RNA was extracted using Qiagen Rneasy Plus Universal Mini kit following manufacturer's instructions (Qiagen). RNA samples were quantified using Qubit 2.0 Fluorometer (Life Technologies) and RNA integrity was checked using Agilent TapeStation 4200 (Agilent Technologies).

RNA sequencing libraries were prepared using the NEBNext Ultra II RNA Library Prep Kit for Illumina using manufacturer's instructions (NEB). Briefly, mRNAs were initially enriched with Oligod(T) beads. Enriched mRNAs were fragmented for 15 minutes at 94 °C. First strand and second strand cDNA were subsequently synthesized. cDNA fragments were end repaired and adenylated at 3'ends, and universal adapters were ligated to cDNA fragments, followed by index addition and library enrichment by PCR with limited cycles.

The sequencing library was validated on the Agilent TapeStation (Agilent Technologies) and quantified by using Qubit 2.0 Fluorometer (Invitrogen) as well as by quantitative PCR (KAPA Biosystems). The sequencing libraries were multiplexed and clustered onto a flowcell on the Illumina NovaSeq instrument according to manufacturer's instructions. The samples were sequenced using a 2x150bp Paired End (PE) configuration. Image analysis and base calling were conducted by the NovaSeq Control Software (NCS). Raw sequence data (.bcl files) generated from Illumina NovaSeq was converted into fastq files and de-multiplexed using Illumina bcl2fastq 2.20 software. One mismatch was allowed for index sequence identification.

**DNaseI-Seq Sample Preparation, Sequencing, and Peak Calling**

DNaseI-Seq was performed as previously described[1–3]. Briefly, 8 x 10^5 - 1.9 x 10^6 cells were lysed using 0.02% IGEPAL. Nuclei were collected by centrifugation at 500 g for 5 min, and DNaseI digestion was performed for 4.5 min at 37°C. DNaseI cleavage fragments were size selected by PEG fractionation, fragments were end repaired, and Illumina sequencing libraries were prepared using the ThruPLEX DNA-seq kit. Libraries were sequenced to a typical mean depth of 44M reads. Reads were processed and peaks of DNaseI cleavages (DHSs) were identified using the default ENCODE DCC DNase-DHS pipeline, version 2.1.2, paired-end (ENCPL848KLD) aligning to GRCh38/hg38. The pipeline uses hotspot2 version 2.1.1 to call peaks at an FDR threshold of 5%. Consensus DNaseI peaks were then identified by merging peak calls from all individual experiments. For each merged element, the full width half maximum of combined DNaseI density from the samples that had a peak overlapping the element was used to identify the core element. All peaks that overlapped the core elements by at least 50% were removed and the process was iterated until all original peaks were represented. To recalibrate the FDR after combining multiple datasets, the combined elements were filtered by the combined z-score for the element. An empiric z-score threshold was chosen at which 95% of DHSs above the threshold would be shared between experimental replicates.

**DNaseI-Seq and RNA-Seq Integration Analysis**

The read count matrix was normalized by median normalization on constitutively accessible/sequenced sites, then rounded to integers to accommodate to DESeq[4], which was used to call differential hypersensitive sites (DHSs). Subsequently, the DHSs were annotated by their nearest genes in hg38[5]. DHSs were regarded as significant if adjusted p value < 0.5 and fold change is greater than 1.5. The same filtering criteria was applied to DEGs when the integration with DHSs was performed. Scatter plots were generated by ggplot2[6] in R4.2.0[7] to visualize the relationship between log2(Fold change) from the two assays.

***In Vitro* Doubling Assay**

Cells from individual clones were seeded to 15cm plates at a known quantity. When cells reached 70-80% confluence, they were trypsinized, diluted in media to a concentration of 1-4e$^{6}$ cells/mL for accurate counting, and sampled three times for counting on a Countess II Automated Cell Counter (Invitrogen). The total number of cells in media was recorded, and a portion of the cells were seeded to a 15cm plate at a known quantity. This process was repeated over approximately two weeks to collect multiple time points for this assay. Total cell counts at each time point were normalized to the number of cells seeded at the previous time point to calculate cell doublings, and like clones (e.g. all SAFE clones, all DKO clones) were combined as replicates when plotting. A general least squares (GLS) approach was used to test for differences in temporal trajectories in doubling counts across time points. Accordingly, a general unspecified covariance structure was assumed to account for the potential within-clone correlation across each time-point. Parameters were estimated by the maximum likelihood method, and tests of fixed effects implemented using likelihood ration statistics. Analyses were performed using the “nlme” (v 3.1-166) R package.

**Necropsy, Sample Preparation, and Immunohistochemistry**

Mice were sacrificed on Day 33 post-injection, which was informed by when the Average Radiance of tumors in the fastest growing group measured about 100,000 [p/s/cm¬≤/sr]. Necropsies were performed to retrieve the brain, spine, liver, and adrenal glands of each mouse, which were promptly placed in formalin. The spines were then placed in PBS, rinsed under running tap water for 30 minutes, and suspended in decalcification solution (0.5M EDTA, pH 8.0). Upon sufficient decalcification, the spines were rinsed under running tap water for 30 minutes, fixed in 4% PFA for 8 hours at RT, rinsed again for 30 minutes, and placed in 70% EtOH until paraffin embedding. Brains, livers, and adrenal glands were rinsed and placed in 70% EtOH until paraffin embedding. Paraffin embedded tissues were trimmed and 2 5μM step-sections were collected (100μM apart) from organ (except for adrenal glands, which were sectioned once) and mounted to slides.

For immunostaining of tumor foci, slides were heated at 58° C for 1 hour, loaded into a Leica Bond RX, and sections were dewaxed at 72 °C before being pretreated with EDTA-based epitope retrieval ER2 solution (Leica, catalog no. AR9640) for 20 minutes at 100° C. A mouse monoclonal primary antibody against human mitochondrial antigen (Millipore, catalog no. MAB1273, diluted 1:1200) was applied for 60 minutes. Slides were then incubated with a rabbit anti-mouse linker (Abcam, catalog no. ab133469, diluted 1:1000) for 8 minutes, followed by incubation with Leica Bond Polymer (anti-rabbit HRP; included in the Polymer Refine Detection Kit (Leica, catalog no. DS9800)) for another 8 minutes. Mixed DAB reagent (Polymer Refine Detection Kit) was then incubated for 10 minutes and followed by Hematoxylin (Refine Detection Kit) counterstaining for 10 minutes. After staining, slides were washed in water, dehydrated using an ethanol gradient (70%, 90%, 100%), washed 3 times in HistoClear II (National Diagnostics, catalog no. HS-202), and mounted in Permount (Fisher Scientific, catalog no. SP15). Slides were then scanned on a Pannoramic 250 Scanner (3DHistech, Budapest, Hungary) using a 20x/0.8NA objective.
